# Multiplexing Light-Inducible Recombinases to Control Cell Fate, Boolean Logic, and Cell Patterning in Mammalian Cells

**DOI:** 10.1101/2025.02.05.636709

**Authors:** Cristina Tous, Ian S. Kinstlinger, Maya E. L. Rice, Jenny Deng, Wilson W. Wong

## Abstract

Light-inducible regulatory proteins are powerful tools to interrogate fundamental mechanisms driving cellular behavior. In particular, genetically encoded photosensory domains fused to split proteins can tightly modulate protein activity and gene expression. While light-inducible split protein systems have performed well individually, few multichromatic and orthogonal gene regulation systems exist in mammalian cells. The design space for multichromatic circuits is limited by the small number of orthogonally addressable optogenetic switches and the types of effectors that can be actuated by them. We developed a library of red light-inducible recombinases and directed patterned myogenesis in a mesenchymal fibroblast-like cell line. To address the limited number of light-inducible domains (LIDs) responding to unique excitation spectra, we multiplexed light-inducible recombinases with our ‘Boolean Logic and Arithmetic through DNA Excision’ (BLADE) platform. Multiplexed optogenetic tools will be transformative for understanding the role of multiple interacting genes and their spatial context in endogenous signaling networks.

## Introduction

The ability to optogenetically control transgene expression has revolutionized our ability to understand cellular signaling. Light can rapidly perturb a protein with high spatiotemporal control(1,2), which is invaluable for mapping the effect of specific genetic regulation onto downstream cell state changes(1,3,4). For example, optical stimulation of channelrhodopsin-2 has been used to delineate sleep state transitions(5), motor cortex function(6), and fear memory recall(7). In comparison to light, chemical inducers traditionally used to control genetic circuits diffuse freely, thwarting the possibility to pattern gene expression(8). While there are many examples of optogenetics transiently modulating cells(9,10), there is a lack of tools to permanently perturb a cell, especially beyond the more heavily explored blue wavelengths. Biological processes are governed by a network of genes dynamic in both space and time, necessitating multi-wavelength control. In the context of skeletal muscle models in mice, fibrosis is impacted by the spatiotemporal expression of various genes (Vim, Fn1, ThbS4), as is regeneration (MyoD, Myl4, Hsp2, Sparc) and calcification (Bgn, Ctsk, Spp1)(11). Multi-wavelength controllable gene circuits will ultimately pave the way for more precise disease and regeneration models critical to understanding the molecular mechanisms underlying cell signaling in those contexts.

Optogenetics takes advantage of photosensitive proteins that can be genetically encoded to offer a layer of light-inducible regulation. Pioneers of the field leveraged opsins, which have traditionally controlled ion flow in neurons, to elicit desired biological responses(1,12,13). A complementary approach involves engineering pairs of dimerization-based optogenetic switches to control the activity of gene editing tools(2,14). Focusing on the latter approach, investigators have developed a vast collection of blue light-dimerization tools utilizing LOV domains(15,16) and cryptochromes(17,18) which are responsive to blue light(19). In addition to blue light domains, investigators have identified and modified red light domains leveraging the red light specific affinity of Phytochromes(20). Red light penetrates tissues deeper than blue light because longer wavelengths result in less scattering and absorption than shorter wavelengths(20,21). By splitting a protein and fusing light responsive domains, researchers have created many unique switches using recombinases(22,23), transcription factors(21,24–26), and Cas proteins(27,28).

Currently, there is an untapped opportunity to multiplex wavelengths, where multiple cell perturbations are controlled independently by light. Orthogonally controlled tools would open the possibility of creating logic-encoded gene circuits and increase the complexity exerted over synthetic circuits. Previous studies have established the possibility of orthogonally controlling genes in the same cell. However, they have not demonstrated Boolean encoded logic, multichromatic patterning, or permanent control over gene outputs.

Two published multichromatic systems multiplexed three light-inducible transcription factors to control the expression of three independent outputs. In the first study, two heterodimerizing transcription factors responsive to UV light (311 nm) and red light (660 nm) were multiplexed with a homodimerizing blue light-inducible (465 nm) transcription factor(29). More recently, a bidirectional, cyanobacteriochrome-based light-inducible dimer (BICYCL) system was engineered to respond to red (660 nm) and green (525 nm) light(30). In addition to developing binders to Amg2, this study multiplexed red, green, and blue (455 nm) heterodimerizing transcription factors. Another innovative multichromatic system was developed using blue (450 nm) light-inducible optoAKT1 and red/far-red (660 nm/ 740 nm) light-inducible optoSOS to control orthogonal cellular signaling pathways(31).

While these studies demonstrate proof-of-concept multiplexing of light inducible gene regulation tools, they have a few drawbacks. First, they require continuous illumination for sustained perturbation. Second, without incorporating Boolean logic- which is challenging with transcription factors and signaling cascades- those circuits can only control one output per wavelength. Finally, there is an absence of gene patterning in two of the three above-mentioned multichromatic circuits and no instances of orthogonally patterned outputs or logic-encoded outputs. There is an urgent need for more gene regulatory tools that enable precise multiplex and programmable control. Expanding the synthetic biologist’s toolbox to include light-inducible recombinases will broaden our spatiotemporal regulatory control to DNA modification, which we can leverage to perform more complex computations than transcriptional regulation-based systems. There is additionally a need to spatially control gene editing to enable the creation of higher-order tissue functions since spatially organized gene expression is a requirement for more complex tissue function. Together, these advancements will push the boundaries of synthetic biology to augment the complexity of circuits possible to ultimately understand the spatial dynamics of biological systems, and to apply spatial control towards applications in regeneration and cell modeling.

Existing multichromatic circuits take advantage of transcription factors and kinases, which require continuous illumination for sustained perturbation. Permanent optogenetic switches would alleviate the need to continuously illuminate cells to achieve sustained perturbation or gene expression, which may be important for applications in disease(32,33) and cell fate pathway modeling(34,35). Site-specific recombinases (SSRs) are an important class of tools that can permanently delete, insert, or reconfigure parts of DNA with high fidelity(36,37). This is advantageous within optogenetics since a single illumination dose is sufficient to permanently activate or repress gene expression. In comparison, transcription factors transiently modify gene expression and require repeat illumination doses to maintain perturbation, expression, or repression. In contrast, the irreversibility of Cre editing makes it especially important to reduce dark activity. Despite this problem, there is an existing blue light-inducible (470 nm) Cre in vivo model that does not suffer from a prohibitive amount of dark activity(23).

Although light-inducible recombinases have been used under the control of one wavelength(22,23,38), they have yet to be multiplexed. Multiplexing recombinases opens the opportunity to engineer cells to perform computations (for example AND, OR, NOT logic) and execute precise biological functions. Promising multiplexing platforms for recombinases include ConVERGD (Conditional Viral Expression by Ribozyme Guided Degradation)(39), INTRSECT (Intronic Recombinase Sites enabling Combinatorial Targeting)(40), and BLADE (Boolean Logic and Arithmetic through DNA Excision). Of these platforms, BLADE is the only architecture that unlocks the opportunity to have an exponential increase in outputs. With BLADE, orthogonal recombinases execute Boolean logic by performing excision at cognate recognition sites on a single transcriptional unit, which has been leveraged with chemical-inducible recombinases. Instead of having 1:1 inputs to outputs with single reporter circuits, 2 recombinase inputs multiplexed with BLADE yields 4 outputs, and 3 inputs yields 8 outputs. In the previously mentioned multichromatic circuits, the number of independent outputs can never be higher than the number of LID-responsive excitation spectra. Boolean logic is one way to circumvent the challenge of a limited number of orthogonal light-inducible domains. One of the greatest advantages of recombinases is the potential to encode sophisticated logic to control multiple outputs within a cell(41–43).

In this paper, we describe the largest collection of red light-inducible recombinases to date. We present 10 red light-inducible recombinases, and we showcase the potential of these tools by patterning a red light-inducible myogenesis switch that turns on MyoD and converts fibroblasts to myotubes. This collection also opens the opportunity for multiplexing logic-based circuits. We select the highest performing red light-inducible recombinases to test in a pairwise screen with blue light-inducible recombinases to carry out AND gate logic. From this screen, we identify 3 red light Flp and blue light Cre pairs with fold changes over 10. This is the first instance of multiplexing red and blue light-inducible recombinases in mammalian cells. We further test orthogonality with our lab’s previously introduced 2-input-4-output BLADE (Boolean Logic and Arithmetic through DNA Excision) decoder and show that we could reproduce BLADE decoder logic with optical rather than genetic or chemical inputs. Finally, we spatially pattern orthogonal gene outputs within mammalian cells for the first time. We expect the multiplexed optogenetic tools described here will be transformative for understanding the role of multiple interacting genes in endogenous signaling networks.

## Results

### Library of Red Light-Inducible Recombinases

Although many laboratories have designed high performing blue light-inducible recombinases to control gene switches in vitro and in vivo(19,23,44), there exists no corresponding library of red light-inducible recombinases. To design red light-inducible recombinases, we fused 3 previously identified candidate red light responsive dimerization domains (Fig. 1a) to 5 different split Cre recombinases. RedMap is a pair of proteins derived from the *Arabidopsis Thaliana* plant phytochrome that utilizes phycocyanobilin (PCB) as a cofactor to dimerize PhyA (a naturally occurring red/far-red light-responsive photoreceptor) and FHY1 (a shuttle protein that sequesters PhyA to the nucleus)(25). In contrast, MagRed(45) and NanoReD(46) are dimerization systems derived from the bacterial phytochrome in *Deinococcus Radiodurans,* which requires biliverdin, an endogenous cofactor available in mammalian cells. While MagRed includes a full-length DrBphP (a naturally occurring bacterial phytochrome) and Aff6 affibody (de novo engineered binder), NanoReD consists of a truncated DrBphP (histidine kinase removed) and an LDB3 nanobody (de novo engineered binder).

**Figure 1.**
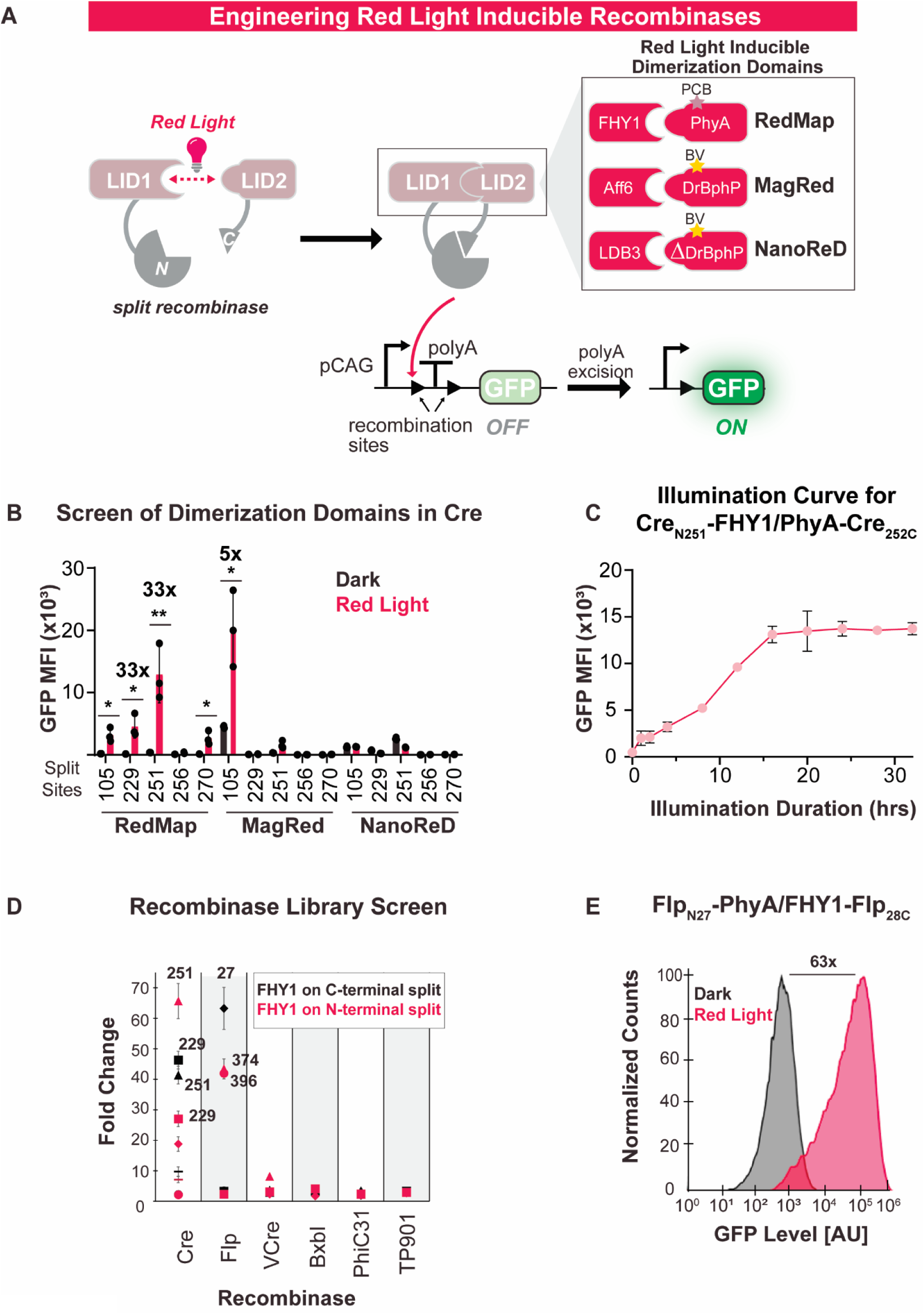
Split site screening to develop a library of high performing red light-inducible recombinases in mammalian cells. (A) Schematic of red light-inducible recombinase circuit. Under red light, recombinase reconstitutes and excises a transcriptional terminator, resulting in a GFP signal. The 3 red light dimerization domains tested are: RedMap, MagRed, and NanoReD with corresponding cofactor indicated by star. (B) Screen of dimerization domains in 5 Cre split sites in HEK cells. Split Cre_N229_-FHY1/PhyA-Cre_230C_ and Cre_N251_-FHY1/PhyA-Cre_252C_ show highest fold changes of 33X. X-axis denotes the final residue of the N-terminal fragment. P-values were calculated by two-tailed unpaired t-test and from left to right: p=0.0108, p=0.0146, p=0.0086, p=0.0161, p=0.0114. (C) Illumination curve indicates optimal light exposure is 20 hours at 0.2 mW/cm². (D) Results of screen fusing RedMap domains to split Cre, Flp, VCre, BxbI, PhiC31, and TP901. Transfection marker gated for top 50% of cells to screen activity of highly expressed split recombinases in HEK cells. (E) Histograms of GFP expression under dark or illuminated conditions for Flp_N27_-PhyA/FHY1-Flp_28C_ show a 63-fold increase. Fold changes were calculated as the ratio of reporter activity with illuminated recombinases over reporter activity with recombinases in the dark. Data represented as mean values ± SD (n=3). All source data is provided in the Source Data file.

We conducted the dimerization screen in HEK cells to identify the highest performing light-inducible domains (LIDs). We transfected 2 split halves of a recombinase fused to dimerization domains in addition to a GFP reporter and a transfection marker. Split sites were chosen based on previous evidence of inducibility in other chemical-inducible recombinases (22). In the presence of red light (47) (using the optoplate shown in Fig. S1), inducible recombinases reconstitute to excise a transcriptional terminator and this results in GFP expression (Fig. 1a). All split Cre pairs in Figure 1b have the photoreceptor fused to the C-terminus (denoted as PhyA-Cre_[Split Site]-C_ for RedMap) and the binder protein fused to the N-terminal fragment of Cre (denoted as Cre _N-[Split Site]_-FHY1 for RedMap).

In this screen, Cre fused to RedMap yielded the greatest number of inducible recombinases. In contrast, Cre fused to MagRed produced one inducible split with high leakiness, and Cre fused to NanoReD showed no inducible activity (Fig. 1b). Within the RedMap-inducible Cre splits, 4 out of 5 split sites resulted in increases exceeding 15-fold. Notably, Cre_N229_-FHY1/PhyA-Cre_230C_ and Cre_N251_-FHY1/PhyA-Cre_252C_ resulted in 33X fold change. Cre_N105_-Aff6/DrBphP-Cre_106C_ also dimerized in the presence of red light with a fold change of 5, but leakiness was high, so we selected the RedMap domains for constructing our larger recombinase library (Fig. S2). To test whether there was an orientation preference for each dimerization system, we also fused each domain to the other split Cre piece (Fig. S3). Similar to the first orientation screened, Cre fused to RedMap showed the highest fold changes of inducibility from light to dark.

We optimized reporter activation of Cre fused to RedMap through transfection of Cre_N251_-FHY1/PhyA-Cre_252C_ in HEK cells. To increase the GFP signal, we illuminated transfected HEK cells at 660 nm from 1 hour to 32 hours at 0.2 mW/cm^2^, and the highest fold change in reporter expression occurred at 20 hours (Fig. 1c). We additionally titrated intensity and pulse duration of red light for 20 hours (Fig. S4a), and intensity for 1 hour (Fig. S4b). We also conducted a PCB dose curve that revealed 100 µM PCB enabled the highest fold change in reporter expression (Fig. S5).

After conducting a screen of LIDs with split Cre, we also fused RedMap domains to split Flp, VCre, BxbI, PhiC31, and TP901 recombinases (Fig. 1d). We hypothesized that split sites, which exhibited strong inducibility when fused to chemical-inducible domains (CIDs) would likewise be effective with LIDs. Ten red light-inducible recombinases exhibited a fold change over 8 (raw MFI values are shown in Fig. S6a-g). The top-performing Flp recombinase was Flp_N27_-PhyA/FHY1-Flp_28C_, which showed a fold change of 63 (Fig. 1e). We also conducted a head-to-head comparison of our top performing red light-inducible Flp with a previously published split Cre using MagRed(38). In our hands, Flp_N27_-PhyA/FHY1-Flp_28C_ had a fold change of 42. Meanwhile, Cre_N104_-Aff6/DrBphP-Cre_106C_) had a fold change of 4.5 (Fig. S7). In our screen of RedMap fused to split recombinases, there was little orientation preference for inducible split sites. 6 inducible splits had the FHY1 domain on the N-terminus, while 4 had the FHY1 domain on the C-terminus (Fig. S8a). We additionally quantified the percentage of split sites that remained inducible when the previously studied CIDs were changed to RedMap domains. 60% of the Cre inducible split sites remained inducible after being fused to RedMap domains instead of CIDs, but this percentage dropped to 37.5% for Flp and 16.66% for VCre (Fig. S8b).

### Red Light-Inducible Myogenesis Switch in C3H10T1/2 cells

An optogenetic recombinase would be particularly useful in cell fate patterning. We leveraged red light-inducible recombinases in C3H10T1/2 mouse fibroblast cells, which are commonly used to study cell fate decisions(48,49). C3H10T1/2 cells can differentiate into myotubes through overexpression of the master transcription factor MyoD(50–52). Upon MyoD induction by red light, our engineered C3H10T1/2 cells are expected to form myotubes, which are multinucleated, elongated cells (Fig. 2a).

**Figure 2.**
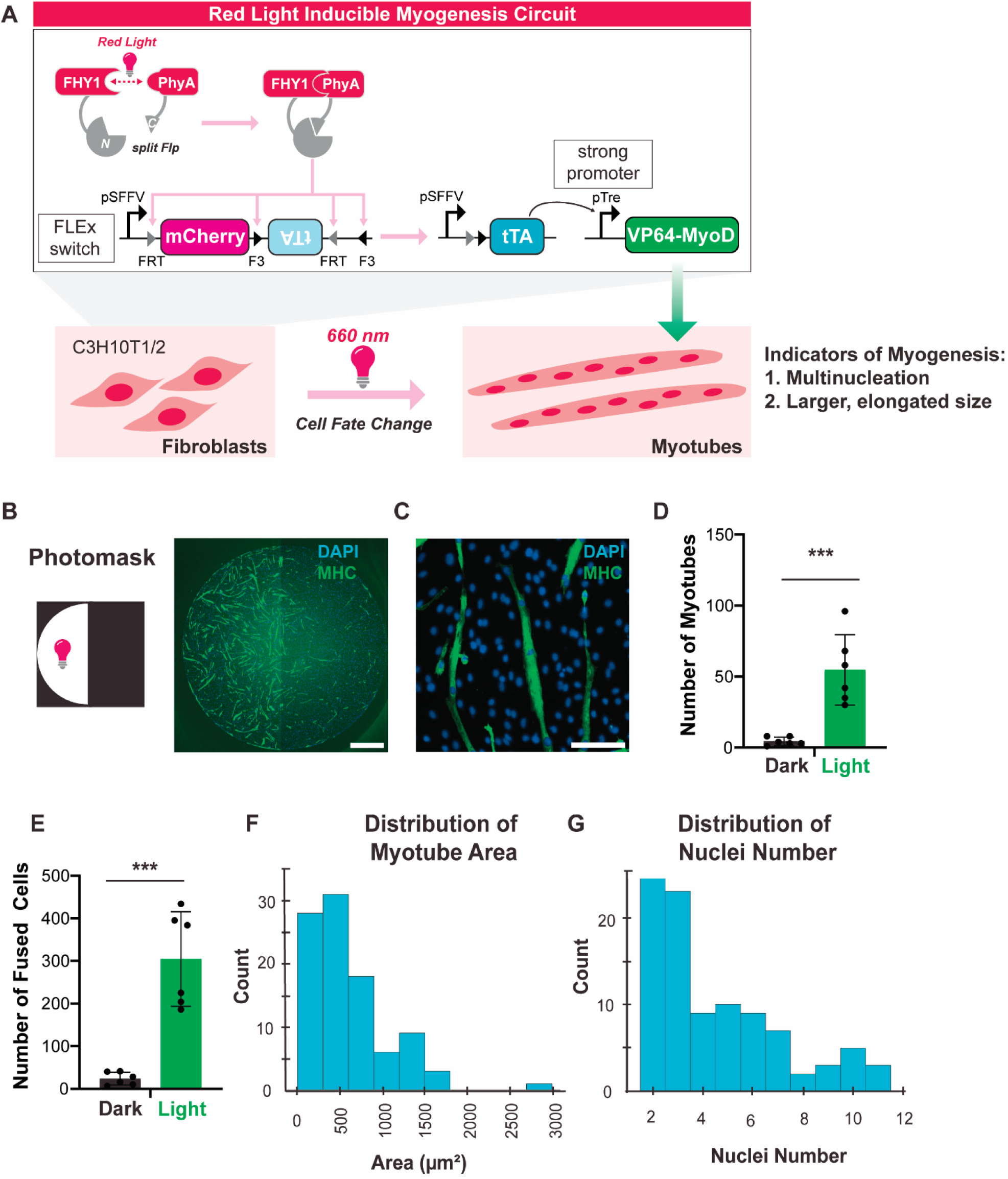
Patterned red light-inducible myogenesis switch in C3H10T1/2 cells. (A) Illustration of myogenesis circuit transduced into C3H10T1/2 cells. Transduced Flp reconstitutes under red light illumination and inverts the sequence encoding tTA. tTA then binds upstream of VP64-MyoD to induce myogenesis. Engineered C3H10T1/2 mouse fibroblasts differentiate into myotubes upon red light illumination due to increased MyoD expression. (B) C3H10T1/2 cells transduced with tTA FLEx switch, inducible MyoD, and Flp_N396_-FHY1/PhyA-Flp_397C_ (denoted as C3H_tTA_ cells). Representative image after illumination through a photomask. Nuclei were stained with DAPI (blue) and myotubes with anti-MHC (green). Scale bar is 1 mm. (C) Higher magnification image of C3H_tTA_ cells. Scale bar is 0.25 mm (D) Overall number of myotubes counted for C3H_tTA_ cells on the left (illuminated) vs right (dark) side of photomask illuminated wells. P-value was calculated as a two-tailed unpaired t-test, p=0.0006. Data represented as mean values ± SD (n=6). (E) Overall number of fused cells counted for C3H_tTA_ cells on the left (illuminated) vs right (dark) side of the well. P-value was calculated as a two-tailed unpaired t-test, p=0.0001. Data represented as mean values ± SD (n=6). (F) Histogram showing the distribution of myotube area (µm²). (G) Histogram showing the distribution of the number of nuclei. All source data is provided in the Source Data file.

We initially designed a non-amplifying MyoD circuit, in which we transduced VP64-MyoD in a Flip-Excision (FLEx) switch and transfected Flp recombinase into C3H10T1/2 cells (C3H_myod_)(53) (Fig. S9a). After illuminating C3H_myod_ cells for 12 hours, we observed morphological changes under the microscope for transfected wild-type (WT) Flp but not for transfected Flp_N396_-FHY1/PhyA-Flp_397C_. Previous studies have calculated changes in myotube formation with western blots(54) and RT-qPCR of MyoD(51), as well as mean Myosin Heavy Chain (MHC) pixel intensity and MHC area with overlapping DAPI-stained nuclei(52,55). We sought to quantify cell fate change using a method that would also capture the morphological changes to these cells. To quantify cell fate changes, we developed a semi-automated image processing algorithm to quantify myotube number by identifying cells stained for anti-MHC conjugated to Alexa Fluor 488 with more than one nucleus. Our algorithm classifies structures as a myotube within the MHC channel if there are at least two overlapping nuclei (Fig. S10a). Thus, to confirm myotube formation, we immunostained C3H_myod_ cells with anti-MHC and DAPI, and we quantified 157 myotubes formed with WT Flp, and 0 with Flp_N396_-FHY1/PhyA-Flp_397C_ (Fig. S9bc).

Next, we engineered an amplification layer to increase the expression of VP64-MyoD after red light reconstitution of split Flp (Fig. S9d). In the presence of red light, transfected split Flp will excise mCherry and turn on a reverse tetracycline-regulated transactivator (rtTA). Then, rtTA will bind upstream of a Tre promoter to activate the expression of VP64-MyoD. We transduced the rtTa FLEx switch and MyoD plasmid into C3H10T1/2 cells (C3H_rtTa_), and then transfected Flp_N396_-FHY1/PhyA-Flp_397C_. Following 12 hours of red light exposure, we counted an average of 90 myotubes, compared to 8 identified in the dark condition, most likely due to spontaneous reconstitution of recombinases at high concentrations (Fig. S9e-i). We also calculated the mean MHC pixel intensity for C3H_rtTa_ cells in the dark or red light illuminated, showing an increase in MHC under red light (Fig. S11).

To further improve our circuit and remove the necessity of doxycycline induction, we replaced rtTA with tetracycline- controlled transactivator (tTA) in the amplification layer (Fig. 2a). We also transduced Flp_N396_-FHY1/PhyA-Flp_397C_ in addition to the amplification circuit into C3H10T1/2 cells (C3H_tTA_). Using this method, we illuminated C3H_tTA_ cells with increasing red light energy doses and saw a corresponding increase in the number of myotubes (Fig. S12). Furthermore, we were able to induce myogenesis with spatiotemporal control using just 30 minutes (63 mJ) of red light, but 1.5 hours of red light illumination showed the best patterning (Fig. 2bc). We calculated 74 myotubes on the illuminated side and 6 on the dark side (Fig. 2d, S13, S14). Additionally, we calculated 404 fused cells under illumination and 32 in the dark, which is quantified as the overall number of nuclei within all of the identified myotubes (Fig. 2e). Finally, we showed the distribution of myotube area and nuclei number of Fig. 2b to showcase the variety of myotube structures observed under illumination (Fig. 2f,g).

### Multichromatic Boolean Logic

Next, we shifted our focus from exclusively using red light-inducible recombinases to multiplexing red and blue light-inducible recombinases. We conducted a pairwise screen in HEK cells by combining our best performing blue light-inducible recombinases(22) with red light-inducible recombinases from Figure 1d. First, we tested the orthogonality of the RedMap and Magnet dimerization systems and showed that Flp_N396_-FHY1/PhyA-Flp_397C_ only responded to red light (Fig. S15a). Similarly, Cre_N251_-nMag/pMag-Cre_252C_ only responded to blue light (Fig. S15b). After confirming the orthogonality of our LIDs, we employed an AND gate reporter to efficiently identify Cre and Flp pairs that simultaneously activated when illuminated (Fig. 3a).

**Figure 3.**
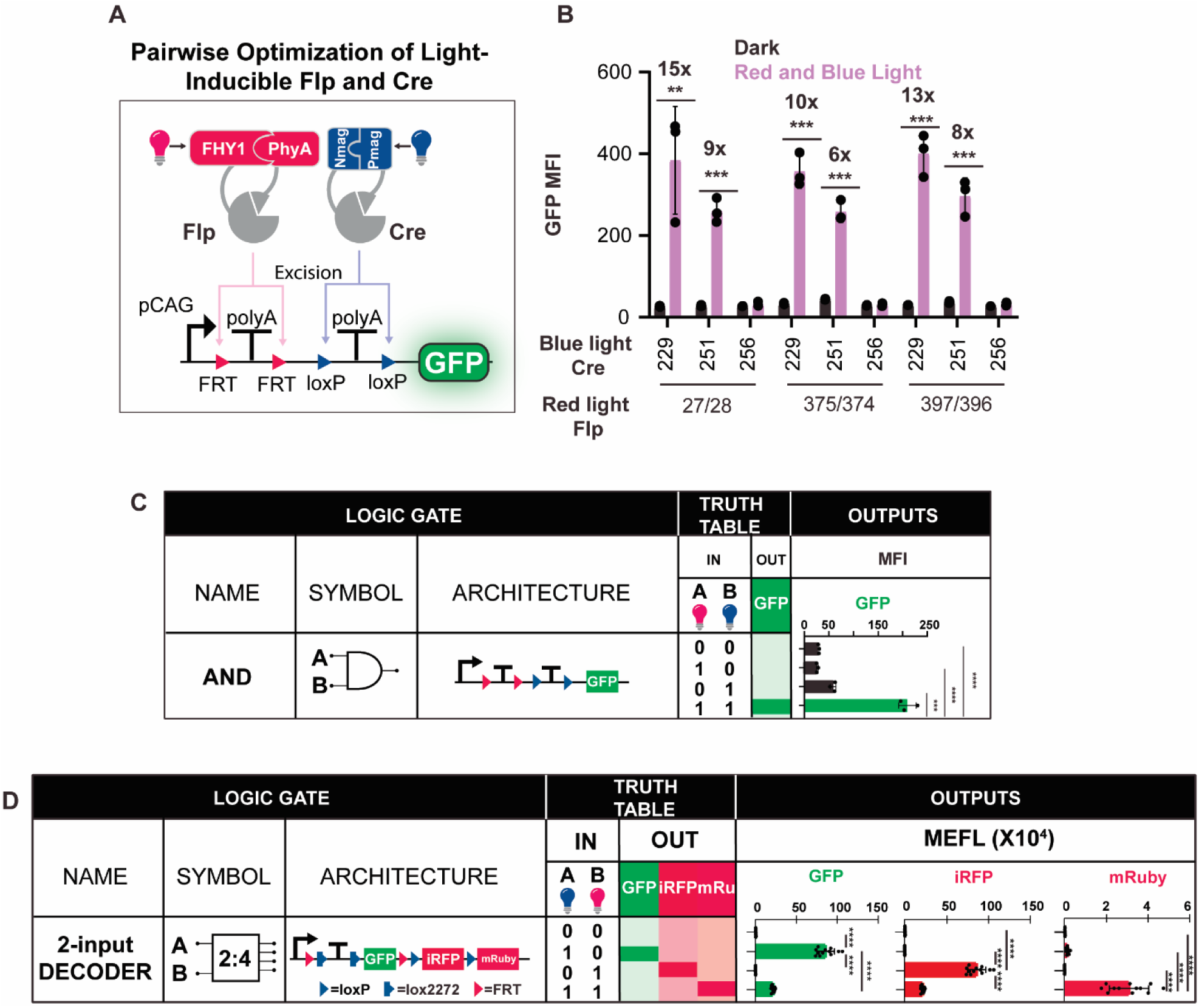
Multiplexing red and blue light-inducible recombinases. (A) Schematic of red and blue light-inducible Cre and Flp circuit transfected into HEK cells. GFP is designed to express only in the presence of both wavelengths. (B) Multiplexed red light-inducible Flp with blue light-inducible Cre. Fold changes were calculated as the ratio of reporter activity with illuminated recombinases over reporter activity with recombinases in the dark. X-axis denotes the final residue of the N-terminal fragment. P-values were calculated as two-tailed unpaired t-tests and from left to right: p=0.0093, p=0.0002, p=0.0002, p=0.0001, p=0.0002, p=0.0005. Data represented as mean values ± SD (n=3) (C) Flp_N396_-FHY1/PhyA-Flp_397C_ and Cre_N251_-nMag/pMag-Cre_252C_ transfected with the AND gate and either illuminated with red light, blue light, both red and blue light, or no light. P-values were calculated as two-tailed unpaired t-tests. For *** p<0.001, **** p<0.0001. (D) Schematic of BLADE 2-input, 4-output decoder transfected in HEK cells. Cre excises lox sites (blue arrows) to express GFP, Flp excises FRT sites (red arrows) to express iRFP, and the presence of both recombinases is designed to express mRuby. Table of light inputs and gene outputs for BLADE transfected with Flp_N396_-FHY1/PhyA-Flp_397C_ and Cre_N251_-nMag/pMag-Cre_252C_. Transfection marker gated for top 40% of cells to screen activity of highly expressed split recombinases in HEK cells. P-values were calculated as two-tailed unpaired t-test where for **** p<0.0001. Data represented as mean equivalent fluorochrome values ± SD (n=12). All source data is provided in the Source Data file.

We tested all possible pairwise combinations of Cre and Flp. We found that the combination of red light-inducible Flp with blue light-inducible Cre yielded six inducible pairs (Fig. 3b), meanwhile red and blue light alone were insufficient to activate the reporter (Fig. 3c). In comparison, blue light-inducible Flp combined with red light-inducible Cre showed no inducible hits (Fig. S16). Following this AND gate screen, we proceeded to test the high performing pairs with our lab’s Boolean Logic and Arithmetic through DNA Excision (BLADE) 2-input decoder.

BLADE is a modular recombinase circuit architecture that takes in Cre and Flp as inputs (Fig. 3d). In circuits with a single reporter per recombinase, recombinases have control over one gene cassette each, but while using BLADE, there is access to an additional, unique third output when both recombinases reconstitute. BLADE was transfected in HEK cells along with Flp_N396_-FHY1/PhyA-Flp_397C_ and Cre_N251_-nMag/pMag-Cre_252C_ (Fig. 3d). We observed that the expected patterns of the reporter were expressed in accordance with the Boolean logic (Fig. 3d, S17). In the dark, there was no fluorescent protein expression because no active recombinases were reconstituted. In the presence of blue light, split Cre reconstituted, which turned on GFP expression. Similarly, in the presence of red light, split Flp was reconstituted and switched on iRFP expression. Finally, in the presence of both wavelengths, mRuby was expressed. GFP and iRFP basal expression occurred under dual red and blue light illumination because Cre and Flp excised DNA unevenly.

### Single and Dual Wavelength Spatial Patterning

To explore spatiotemporal control using these tools, we patterned gene outputs in HEK cells using a single red wavelength and dual wavelengths of red and blue light. Initially, we focused on red light patterning and transfected Flp_N396_-FHY1/PhyA-Flp_397C_ with a GFP reporter into HEK cells. Then, we illuminated cells through a photomask, and we saw considerable GFP activity in the dark regions. We hypothesized a) that red light scattered to the off region to reconstitute split Flp, and b) that basal reconstitution of split recombinases was too high under a CAG promoter. To reduce red light scattering, we plated transfected cells in media containing 0.5 mM Brilliant Blue, an FDA-approved dye that absorbs light in the 660 nm range(56). The addition of Brilliant Blue decreased dark activation in patterned wells by 69% (Fig. S18). We additionally cloned our split recombinases under a doxycycline-inducible promoter to more easily control Flp expression levels (Fig. S19). We saw that 2 µg/ ml doxycycline resulted in the greatest fold change for patterning. After implementing the Brilliant Blue photoabsorber and doxycycline-inducible split Flp, followed by red light illumination, we quantified a steep increase in GFP expression at the photomask boundary (Fig. S20).

Next, we transfected Flp_N396_-FHY1/PhyA-Flp_397C_ and Cre_N229_-nMag/pMag-Cre_230C_ (top performing blue light-inducible Cre) with orthogonal GFP and BFP reporters (Fig. 4a). To pattern half a well with each recombinase reporter, we designed two photomasks which were sequentially attached to the bottom of the plate covering either the right half or top half of each well, and then illuminated through the photomask. With this recombinase pair, we conducted a doxycycline curve from which we chose to induce with .02 µg/ml due to its lower dark activity while preserving high fold change (Fig. S21). Then, in Figure 4bc, we repeated this transfection and after illumination, we observed GFP activated by red light-inducible Flp clearly expressed on the left side, and BFP activated by blue light-inducible Cre was expressed on the right side (Fig. S22). Finally, we transfected Flp_N396_-FHY1/PhyA-Flp_397C_ and Cre_N229_-nMag/pMag-Cre_230C_ with an AND gate reporter to see if we could pattern encoded logic (Fig. 4d). We observed substantial GFP activation by the illumination of red and blue light (Fig. 4ef, S23).

**Figure 4.**
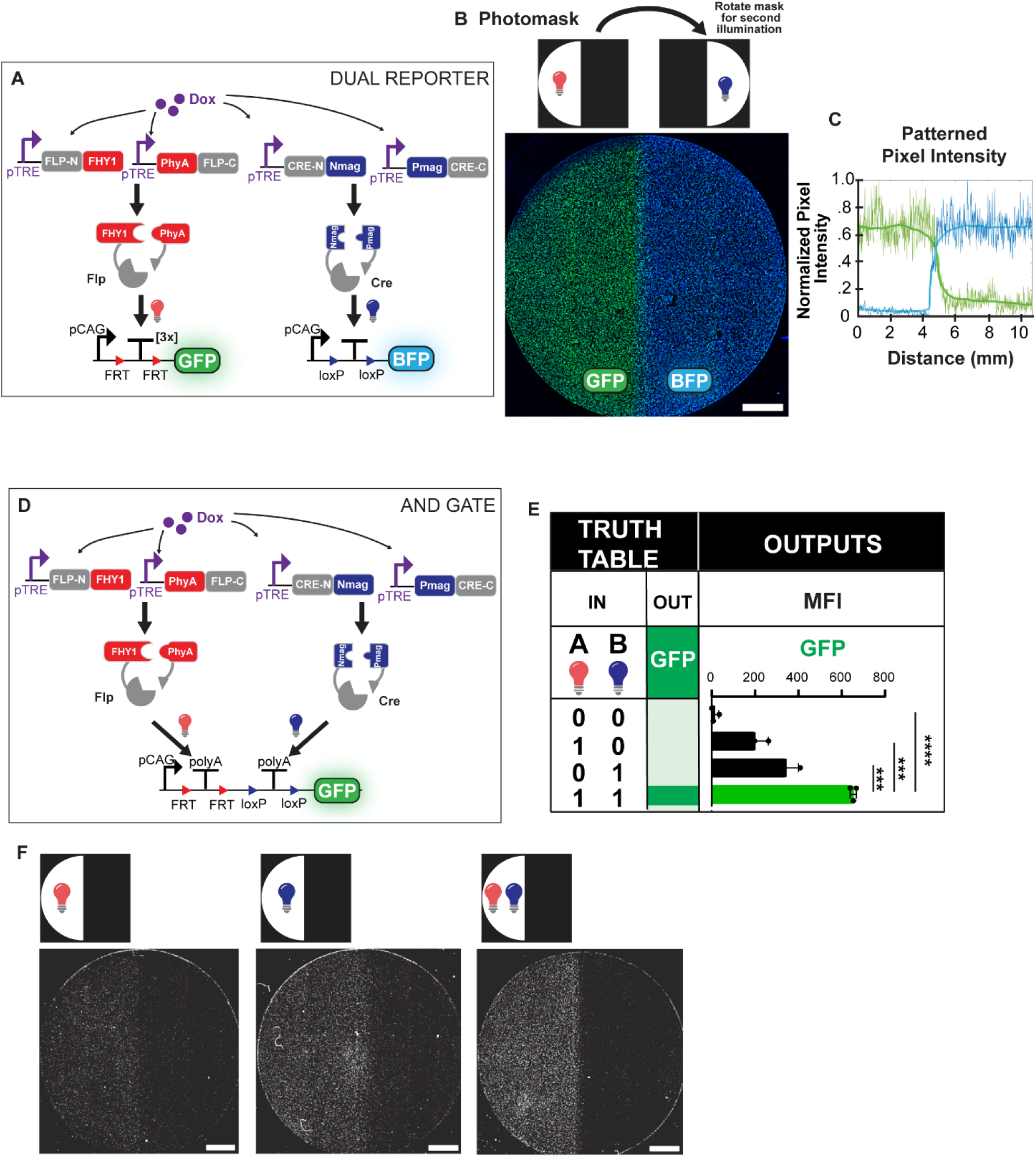
Single and Dual Wavelength Spatial Patterning. (A) Diagram of red light-inducible Flp and blue light-inducible Cre circuits transfected for patterning experiments. (B) Image of photomask patterned HEK cells transfected with Flp_N396_-FHY1/PhyA-Flp_397C_ and Cre_N229_-pMag/nMag-Cre_230C_. Red light-inducible Flp drives GFP on the left half of the well and blue light-inducible Cre drives BFP expression on the right half of the well. (C) Plot of patterned pixel intensity of red light-inducible Flp and blue light-inducible Cre. In blue is the intensity values for Cre and green indicates intensity values for Flp. (D) Diagram of red light-inducible Flp and blue light-inducible Cre circuits transfected with AND gate reporter. (E) Table of light inputs and gene outputs for AND gate transfected with Flp_N396_-FHY1/PhyA-Flp_397C_ and Cre_N229_-pMag/nMag-Cre_230C_ (F) Images of photomask patterned HEK cells transfected with Flp_N396_-FHY1/PhyA-Flp_397C_, Cre_N229_-pMag/nMag-Cre_230C_ and an AND gate reporter. All scale bars 2 mm. All source data is provided in the Source Data file.

## Discussion

In this paper, we tested three different LIDs with split Cre recombinase and found the highest inducibility with RedMap domains. One drawback of RedMap domains is that the photoreceptor PhyA requires the exogenous addition of PCB, compared to MagRed and NanoReD, which do not. Despite this hurdle, in vivo experiments have been done with RedMap-inducible circuits to modulate insulin levels(25). Additionally, PCB can be genetically encoded to overcome its delivery challenges(57).

Furthermore, after conducting a screen of RedMap fused to published chemical-inducible recombinases, we were surprised only 25% of the split sites we tested were inducible. We hypothesized that Cre yielded the most inducible split sites because it has previously performed the most robustly. Our lab’s best performing chemical-inducible Cre and Flp resulted in a fold change of around 275 and 418, while the top performing chemical-inducible TP901 and BxbI recombinases resulted in a lower fold change around 76 and 32(22). We are not sure why our split recombinases have lower performance when fused to RedMap domains. We hypothesize that the larger gene sequence of PhyA (around 1.85 kilobase pairs) may lower protein expression relative to smaller dimerization domains(58,59). Additionally, PhyA is a light labile protein, which could reduce the expression of the split protein compared to chemical-inducible proteins.

One shortcoming of this screen was that our high performing split Flp was not strong enough to induce myogenesis in C3H10T1/2 cells without adding an amplification layer. Developing higher performing red light-inducible recombinases in the future could omit the need for this layer of regulation. Additionally, after testing nine pairs of red light-inducible Cre and blue light-inducible Flp, none showed ON activity with the AND gate reporter. This was most likely due to the lower performance of the blue light-inducible Flp recombinases compared to the blue light-inducible Cre recombinases(22). Finally, our AND gate circuit performance could be further optimized to reduce red and blue light activation of the GFP reporter. These results highlight the importance of developing a collection of high performing recombinases with a large fold change. Our current method employs manual cloning of all subsets of split recombinases. However, exploring all possible splits is not tractable. In the future, it would be beneficial to complement empirical testing with protein structural prediction tools such as AlphaFold while developing a collection of split proteins. Furthermore, the directed evolution of RedMap domains could improve activation kinetics for future applications.

While we and others have demonstrated the utility of individual light-inducible recombinases, here we multiplex red and blue light-inducible recombinases in mammalian cells for the first time. This toolset expands the type of gene regulation possible to encompass Boolean logic not possible with just one wavelength. Future optogenetic recombinase collections should focus on increasing the number of orthogonal light-inducible tools to increase the complexity of Boolean logic that can be done. Traditionally, optogenetic site-specific recombinases have been leveraged for targeted integration or excision of DNA in the development of transgenic animals and in vitro tissue models(60,61). We envision the light-inducible genetic reprogramming of cells will enable mechanistic investigations of cell biology and the development of multicellular engineering living systems. For example, it is well documented that overexpression of transcription factors can direct cell differentiation(51,62). In the future, the fluorescent proteins in our BLADE circuit could be swapped for transcription factors and then integrated into induced pluripotent stem cells (iPSCs) to spatially control cell fate. Sustained gene expression is often needed in various disease and cell fate pathway modeling(63,64). While existing multichromatic circuits require continuous illumination for persistent perturbation, permanently encoded outputs to minimize illumination could reduce toxicity. Conversely, we also recognize that transient methods of gene induction are more likely to reach a lower equilibrium of basal activity, compared to recombinases which have irreversible dark activity that may compound over time.

While optogenetics uniquely uses light to control protein activity and downstream gene expression, complementary fields like biomaterials(65) and 3D bioprinting(66) also strive to control cell signaling and geometry through controlled depots and precise cell deposition. These fields have achieved great advancements in spatial control, but biomaterials and 3D printing alone cannot introduce engineered signaling circuits with sustained signals and genetically encoded logic. Sustained signals are present in a multitude of endogenous cellular pathways, such as control over cell fate(34,35). Meanwhile, genetically-encoded logic increases the number of controllable orthogonal outputs and the complexity of circuits we can design. Multichromatic gene schemes hold the potential to demonstrate sustained signals and genetically encoded logic, although existing publications do not exhibit this. Overall, optogenetics holds tremendous promise in its ability to spatiotemporally control regulatory proteins. This will solve a fundamental problem in regenerative medicine in solving how to orchestrate spatial activation and repression of gene expression within native tissues.

The promise of optogenetics lies in the power to spatially control outputs. Thus, it is crucial to demonstrate spatial control and to design more complex spatial patterns with the help of logic to push the number of gene perturbations possible with optogenetics. We anticipate the tools developed in this work will expand the circuit-encoded logic possible within mammalian cells for pivotal advances in genetically engineering more spatiotemporally precise cellular patterning.

## Materials and Methods

### Mammalian Cell Culture

HEK293FT and C3H10T1/2 cells were cultured at 37 °C with 5% CO_2_. HEK cells were maintained in Dulbecco’s Modified Eagle’s Medium (DMEM) containing 5% fetal bovine serum, 1% penicillin and streptomycin (Corning 3000IC1), 1% L-glutamine (Corning 2500SCI) and 1% sodium pyruvate (Lonza 13115E). C3H10T1/2 cells were maintained in Minimum Essential Medium Eagle supplemented with 10% fetal bovine serum, 2 mM L-glutamine, and 50 UI/ml penicillin and streptomycin. C3H10T1/2 cells were not maintained above passage 15.

### Inducible Split Recombinase and Reporter Plasmid Construction

RedMap domains(25) FHY1 and PhyA were ordered as g-block sequences (Invitrogen) with homologous stretches that overlapped with the Cre backbone for Gibson assembly. The MagRed domains(45) were also ordered as g-blocks (Invitrogen), and the NanoReD domains(21) were a kind gift from Dr. Liangcai Gu. All other plasmids cloned using restriction enzyme digestion and Gibson assembly. All constructs confirmed via a test cut run on gel electrophoresis and then sent for sanger sequencing from Quintara Biosciences. Construct information in Supplementary Table 1.

### Small molecule Preparation

Phycocyanobilin: PCB (SiChem, SC-1800) was dissolved in 100% DMSO to a final 100 mM stock and stored at −20 °C.

Doxycycline: Doxycycline (Gold Biotechnology, D-500-1) was dissolved in water to a final concentration of 2 µg/ml and stored at −20 °C.

### Optoplate Calibration

To calibrate each optoplate, we used optoConfig-96 software(67) to program 14 increasing intensity values between 0 and 4000. For each wavelength, we averaged triplicate power reads and then made a standard curve to calculate the intensity value of each grayscale intensity input on the optoConfig-96.

### HEK Transfection, Illumination, and Flow Cytometry

For flow cytometry experiments, HEK293FT cells were plated in triplicate at 20,000 cells/well in a black-walled 96 well plate and then transfected the next day at 70-80% confluency. In each split red light-recombinase screen, cells were transfected using 0.323 g/L polyethylenimine (PEI) (Polysciences, 23966-2) dissolved in 0.15 M NaCl (4 parts PEI: 25 parts NaCl). After a 10 minute incubation of DNA added to the PEI-NaCl complex, 10 µl transfection volume was dispensed per well. Transfection ratios for red light-inducible screens are detailed in Supplementary Table 2. 2 hours after transfection, PCB was added to cells at a final concentration of 100 µM. 24 hours after transfection, cells were illuminated for 20 hours using an optoplate (47), which was programmed using optoConfig96 software(67) (Fig. S1). 48 hours after transfection, HEK cells were analyzed using a Thermo Fisher Attune Nxt cytometer.

Transfection for the multiplexing screen with AND gate reporter was conducted in triplicate according to ratios detailed in Supplementary Table 3. BLADE experiments were transfected in 6 replicates according to DNA ratios in Supplementary Table 4. Then, 2 wells were pooled together for flow cytometry analysis. Both the AND gate and BLADE transfections were induced with PCB 2 hours post-transfection. Then, 2 hours after drug induction, illumination for the AND gate and BLADE experiments was initiated. For both Boolean logic gates, red light was pulsed for 5 min on, 25 min off for 44 hours at 0.35 mW/cm^2^. Blue light was illuminated 1 day after transfection and was pulsed for 5 min on, 25 min off for 20 hours at 2.5 mW/cm^2^. BLADE transfection fluorescence channel measurements were converted to Molecules of Equivalent Fluorescein (MEFL) units using SPHERO Calibration beads (Spherotech, RCP-30-5A).

### HEK Transfection, Illumination, and Microscopy

HEK293FT cells were seeded into a T-75 at 4.8 million cells per flask. The following day, HEK cells were transfected at around 70-80% confluency with 19,500 ng total of circuit components and then wrapped in aluminum foil. Transfection ratios are detailed in Supplementary Tables 5-7. The next day, the cells were passaged in the dark and reseeded into a black-walled 24 well plate (Ibidi, 82406) at 1 million cells/well. For red light patterning, cells were seeded in media with 0.22 µm filtered 0.5 mM Brilliant Blue (Spectrum, FD110-100GM), 100 µM PCB and 2 µg/ml doxycycline. For red and blue light patterning, cells were seeded in media with 0.22 µm filtered 0.5 mM Brilliant Blue and 2 mM tartrazine (Chem-Impex, 22937), 100 µM PCB and 0.02 µg/ml doxycycline. 5 hours later, cells were illuminated with an Acetal (McMaster-Carr, 8492K212) based photomask taped underneath the culture plate. 0.125 inch Acetal was laser-cut with Epilog Engraver using a speed of 30 and a power of 100 in raster mode, then in vector mode a speed of 30 and maximum power and frequency. Illumination for patterning was done with an RGB projector (EKB Technologies, DPM-E4500MKIIRGBHPCR-OX). Red light illumination was 3 min with the current set to 105. For red and blue light illumination with individual recombinase reporters, red light illumination was the same as single red wavelength patterning, and blue light illumination was 5 min with the current set to 50. For red and blue light illumination with the AND gate, red and blue light illumination was 10 min with current set to 105 and 50. Cell imaging was done the next day on a BioTek Cytation 5 imaging reader. Pixel intensity was measured over each well in Fiji. Raw pixel intensity values were smoothed using the moving average function in MATLAB.

### C3H10T1/2 Transfection

C3H10T1/2 cells were plated at 5,000 cells per well in a black-walled 96 well plate and transfected the following day with Lipofectamine 3000 according to the manufacturer’s protocol (100 DNA ng/well).

### Lentivirus Production for C3H10T1/2 cells

HEK cells were initially plated at 4.8 million cells in a T-75. The next day, at around 80% confluency, HEK cells were transfected with 9.43 µg viral packaging DNA (6.43 µg pDelta, 2.14 µg pVsvg, and 0.86 µg pAdv) and 2.1 µg plasmid of interest (concentrated at more than 1 µg/µl) with 0.16 fraction PEI in 0.15 M NaCl. Media was changed the following day, and for the next 2 days, the virus was collected and stored at 4 °C. On day 5, the virus was spun down at 1200×g for 10 min at 4 °C to remove cellular debris. The supernatant was then transferred to a new conical and concentrated with centrifugal filters (EMD Millipore, UFC801024).

### C3H10T1/2 Cell Transduction and Myogenesis Experiment

C3H10T1/2 cells were plated at 100,000 cells/ well in 2.5 mls in a 6-well plate. The next day, all virus was added dropwise to C3H10T1/2 cells along with 1 µl 10 mg/ml polybrene (EMD Millipore, TR-1003-G). C3H_myod_ cells were transduced with MyoD non-amplifying circuit. C3H_rtTA_ cells were transduced with rtTA FLEx switch and MyoD plasmid in a 1:1 ratio. C3H_tTA_ cells were transduced with tTA FLEx switch, MyoD plasmid, and Flp_N396_-FHY1/PhyA-Flp_397C_ in a 1:1:1 ratio. Cells were passaged for the next two days, and on the second day, cell transduction efficiency was confirmed via flow cytometry. Transduced C3H10T1/2 cells were plated in a black-walled 96 well plate at 5,000 cells/well and then C3H_myod_ and C3H_rtTA_ cells were transfected with Flp_N396_-FHY1/PhyA-Flp_397C_ the following day with Lipofectamine 3000. 2 hours post transfection, cells were induced 100 µM PCB. C3H_rtTA_ cells were also induced with a final concentration of 2 µg/ml doxycycline. Cells were illuminated at 660 nm the next day for 12 hours using our optoplate for C3H_myod_ and C3H_rtTA_ cells. C3H_tTA_ cells were illuminated for 1.5 hours using our projector. The day following illumination, cell media was changed and C3H_tTA_ cells were fixed 2 days later. For C3H_myod_ and C3H_rtTA_ cells, cells were fixed one week after transfection and cell media was changed every 2 days (doxycycline was added every 2 days for C3H_rtTA_ cells).

### C3H10T1/2 Cell Immunostaining

Cells were initially fixed using 4% paraformaldehyde. After fixation, cells were treated with permeabilization solution (0.5% Triton X-100 in PBS) for 10 min and then blocking solution (1% BSA in PBS) for 30 min. Next, cells were stained with anti-Myosin Heavy Chain conjugated to Alexa Fluor 488 (Developmental Studies Hybridoma Bank, MF-20) and diluted in washing solution (.01% Triton X-100 in PBS) at 4 °C overnight. The next morning, cells were washed twice and then incubated at room temperature with anti-mouse IgG Alexa Fluor 488 (Thermo Fisher Scientific, A-21202) diluted in wash solution for 1 hour. Afterwards, the cells were washed twice with wash solution and finally were counterstained with 1 ng/ml of DAPI for 30 min. Finally, the cells were washed once with PBS and stored in PBS at 4 °C.

### Microscopy

Fluorescence microscopy was performed using the BioTek Cytation 5 Cell Imager. A 4x objective was used to collect images at 37 °C, 5% CO_2_ and 90% humidity. For imaging of mCherry, a 554 LED cube was paired with an Ex556/Em600 filter cube. To image GFP and Alexa Fluor 488, a 465 LED cube was paired with an Ex469/Em525 filter cube, and to measure BFP and DAPI, a 365 LED cube was paired with an Ex377/Em447 filter cube.

Confocal microscopy was performed using the Nikon Ti2-E with Andor Dragonfly 505 System. A 20x objective was used to collect images at room temperature. To image anti-MHC conjugated to Alexa Fluor 488, a 465 LED cube was paired with an Ex469/Em525 filter cube, and to measure DAPI, a 365 LED cube was paired with an Ex377/Em447 filter cube.

### Image Processing Software

We developed a semi-automated script to segment and characterize myotube structures in imaging data, which we implemented in MATLAB. The inputs to the script are fluorescence images of myotubes (i.e., cells stained against myosin heavy chain (MHC)) and of nuclei, and the outputs are a series of measurements including a number of myotubes, geometric properties of myotubes, and number of nuclei per myotube. Within the script, the input images are binarized with a user-defined threshold and pre-processed to remove noise. Connected sets of pixels within the MHC channel are classified as candidate myotubes or non-myotubes based on empirically determined thresholds for area and eccentricity. The nuclei channel is used to determine the number of nuclei associated with each candidate myotube, and candidates with a single nucleus are discounted. Because of the complex morphologies of myotubes and the challenges associated with completely automated segmentation, we included an interactive quality control step where the user can manually discount myotubes that were incorrectly identified.

### Flow Cytometry

For red light domain and split site screening experiments, HEK cells were passaged and resuspended in HEK media. All samples were gated for viability on a flow cytometer using the forward scatter (FSC) and side scatter (SSC). Then, transfected populations were gated for a transfection marker and subsequently measured for GFP MFI within that population. No fewer than 10,000 cells were looked at for each sample. Fluorescence data was collected for iRFP (excitation laser: 638 nm, emission: 720/30 nm), GFP (excitation laser: 488 nm, emission: 510/10 nm), mCherry/mRuby (excitation laser: 561 nm, emission: 620/15 nm), LssmOrange (excitation laser: 405 nm, emission: 603/48 nm). Flow cytometry data was then analyzed using FloJo.

### Statistical Methods

All transient experimental analysis via flow cytometry included at least 3 separate cell culture transfections. Fluorescence intensity values were then averaged, and standard deviation was reported. All statistical analysis was performed within Prism 8.

