## Supplement for "Multiplexing Light-Inducible Recombinases to Control Cell Fate, Boolean Logic, and Cell Patterning in Mammalian Cells"

Cristina Tous *et al.*

**This PDF file includes:**

Figures S1 to S23

Tables S1 to S7

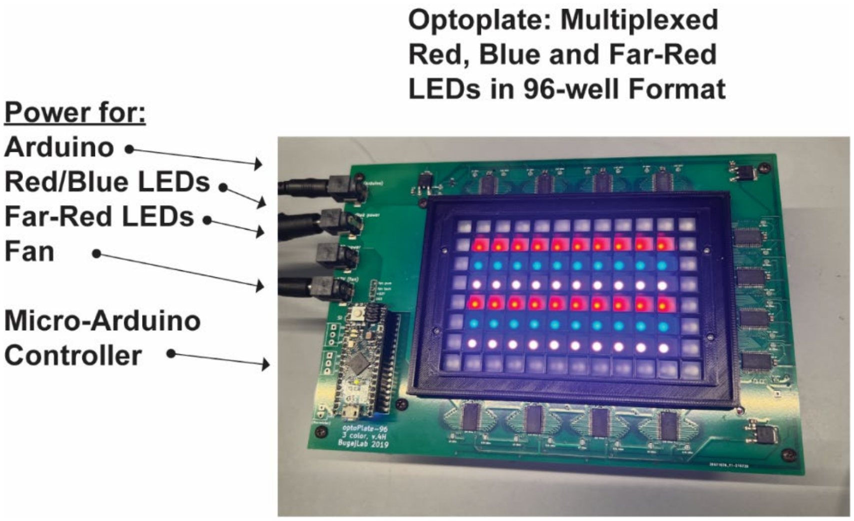


**Supplementary Figure 1. OptoPlate used for illumination**

The optoplate (designed by Lucasz Bugaj and Wendell Lim) was a convenient platform for studying optogenetic circuit perturbation in a high-throughput manner. The optoplate was utilized in Figures 1-3 to illuminate red and blue light.

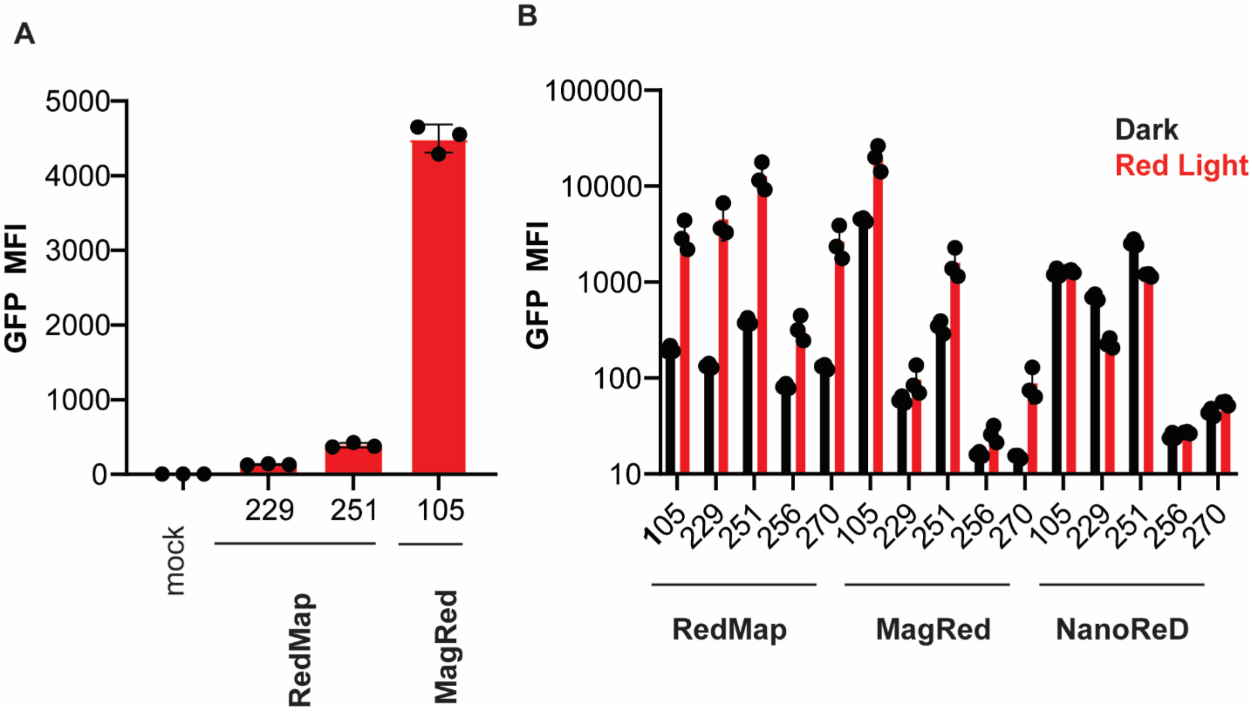


**Supplementary Figure 2. Basal activity of red light-inducible Cre recombinase**

(A) Dark activity in HEK cells of a mock control, $Cre_{N229}$-FHY1/PhyA-$Cre_{230C}$, $Cre_{N251}$-FHY1/PhyA-$Cre_{252C}$, and $Cre_{N105}$-Aff6/DrBphP-$Cre_{106C}$. (B) The screen of dimerization domains in 5 Cre split sites (same graph as Fig. 1b) is shown on a log scale to highlight basal activity. Data represented as mean values ± SD (n=3).

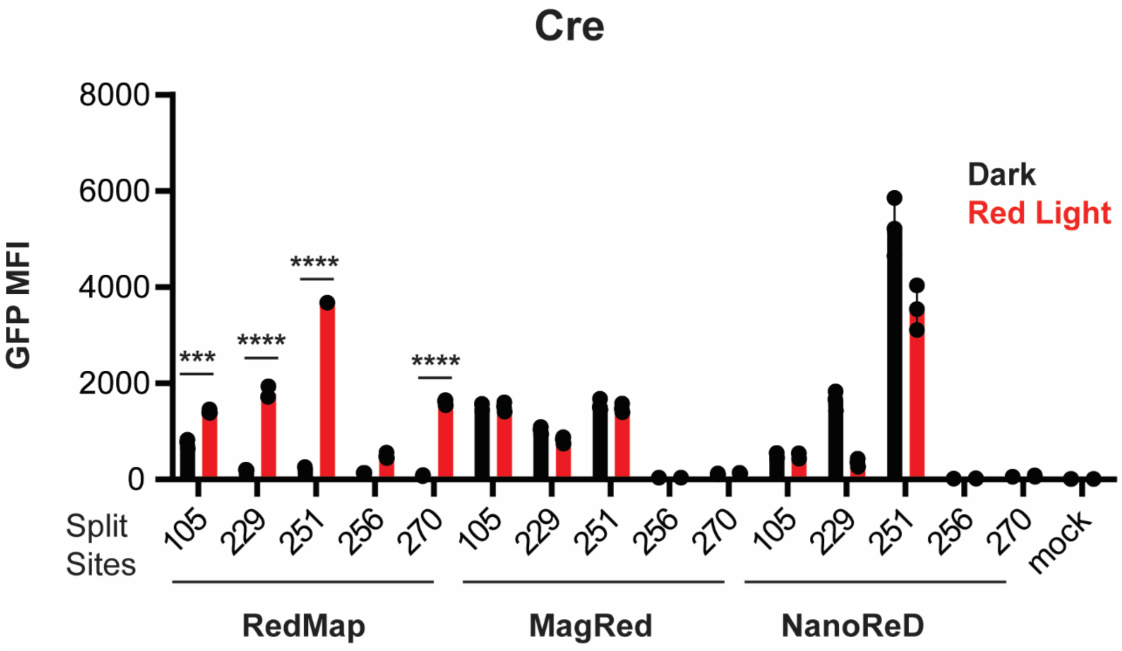


**Supplementary Figure 3. Red light dimerization screen with photoreceptor fused to N-terminus**

RedMap, MagRed, and NanoReD were fused to split Cre in the opposite orientation compared to Figure 1b. PhyA from the RedMap domains and DrBphP from the MagRed and NanoReD domains were fused to the N-terminal fragment. The results show that only split proteins fused to the RedMap domains yielded functionally inducible recombinase activity. X-axis shows split sites on N-terminus of split Cre.  P-values were calculated by two-tailed unpaired t-test, for *** $p<0.001$ and for ****$p<0.0001$. Data represented as mean values ± SD (n=3).

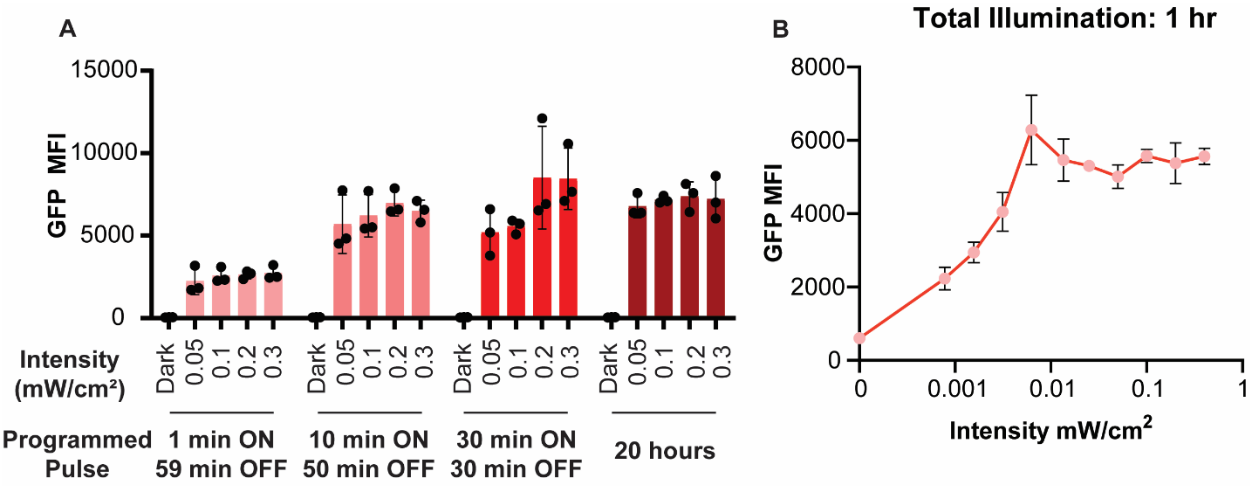


**Supplementary Figure 4. Characterization of red light inducibility**

(A) $Cre_{N251}$-FHY1/PhyA-$Cre_{252C}$ transfected in HEK cells illuminated with different intensities and pulse durations over 20 hours. (B) $Flp_{N396}$-FHY1/PhyA-$Flp_{397C}$ transfected in HEK cells illuminated with different intensities over 1 hour. Data represented as mean values ± SD (n=3).

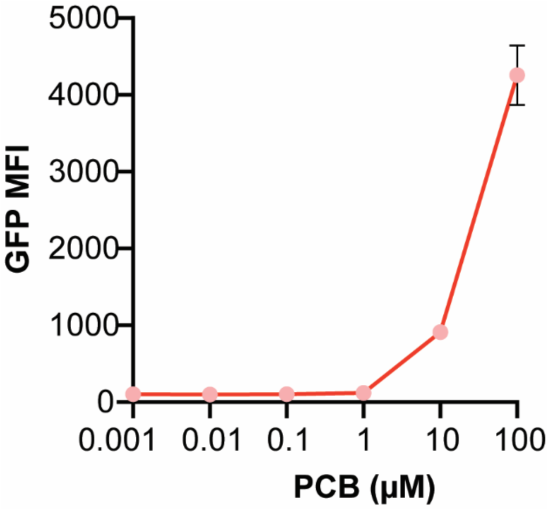


**Supplementary Figure 5. Phycocyanobilin dose curve**

The dose curve indicates that 100 µM phycocyanobilin is the optimal concentration to induce $Cre_{N251}$-FHY1/PhyA-$Cre_{252C}$ transfected in HEK cells. Dose curve did not go to 1000 µM due to insolubility. Data represented as mean values ± SD (n=3).

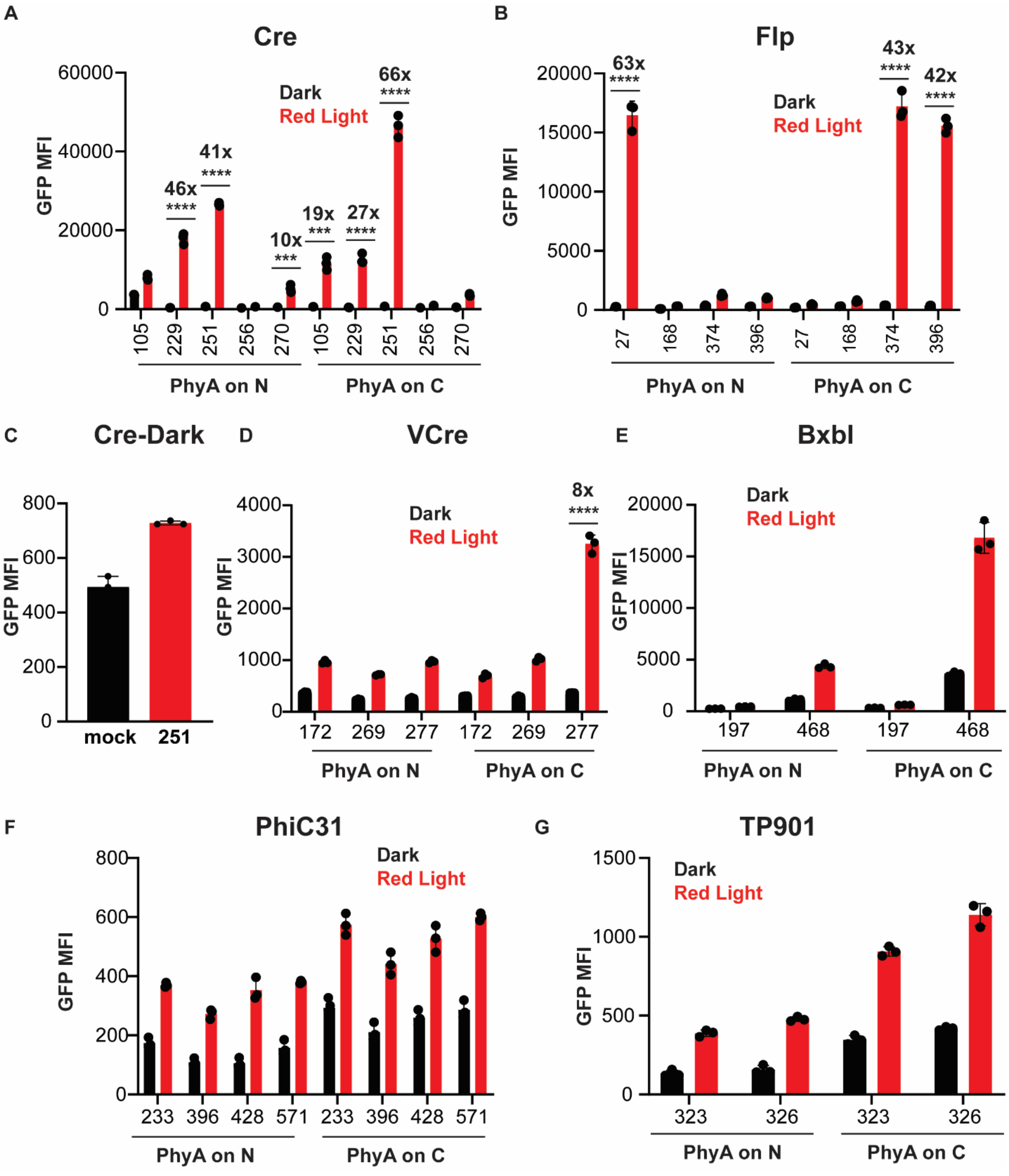


**Supplementary Figure 6. Raw MFIs from multidimensional recombinase screen**

Screens of split recombinases in (A) Cre, (B) Flp, (C) dark activity of $Cre_{N251}$-FHY1/PhyA-$Cre_{252C}$ compared to mock control. Split recombinase screens in (D) VCre, (E) BxbI, (F) PhiC31, and (G) TP901. Transfection done in HEK cells and reconstituted recombinases turn on GFP reporter expression. P-values were calculated by two-tailed unpaired t-test. For *** p<0.001 and for ****p<0.0001. Data represented as mean values ± SD (n=3).

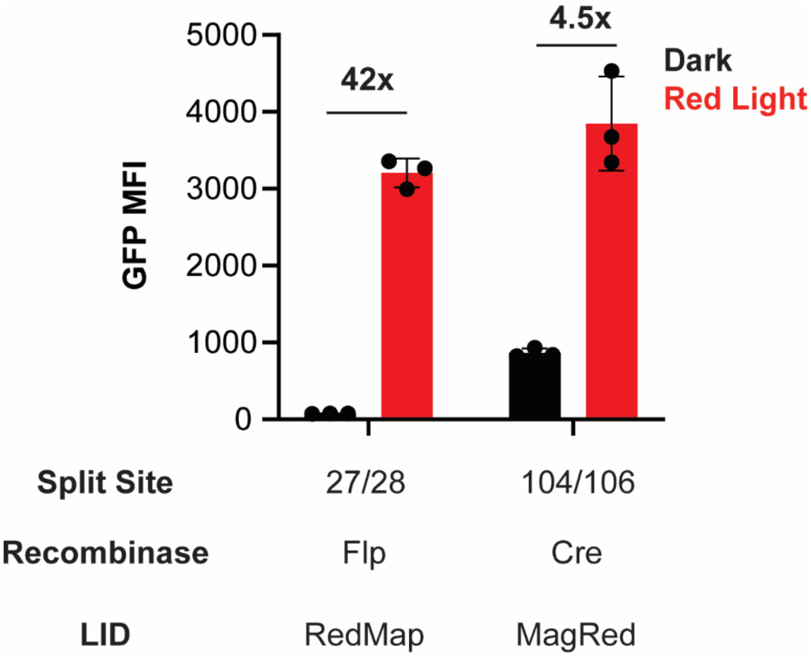


**Supplementary Figure 7. Red light-inducible recombinase comparison**

Graph of $Flp_{N27}$-PhyA/FHY1-$Flp_{28C}$ and $Cre_{N104}$-Aff6/$Cre_{106C}$-DrBphP transfected with GFP reporter in HEK cells. Data represented as mean values ± SD (n=3).

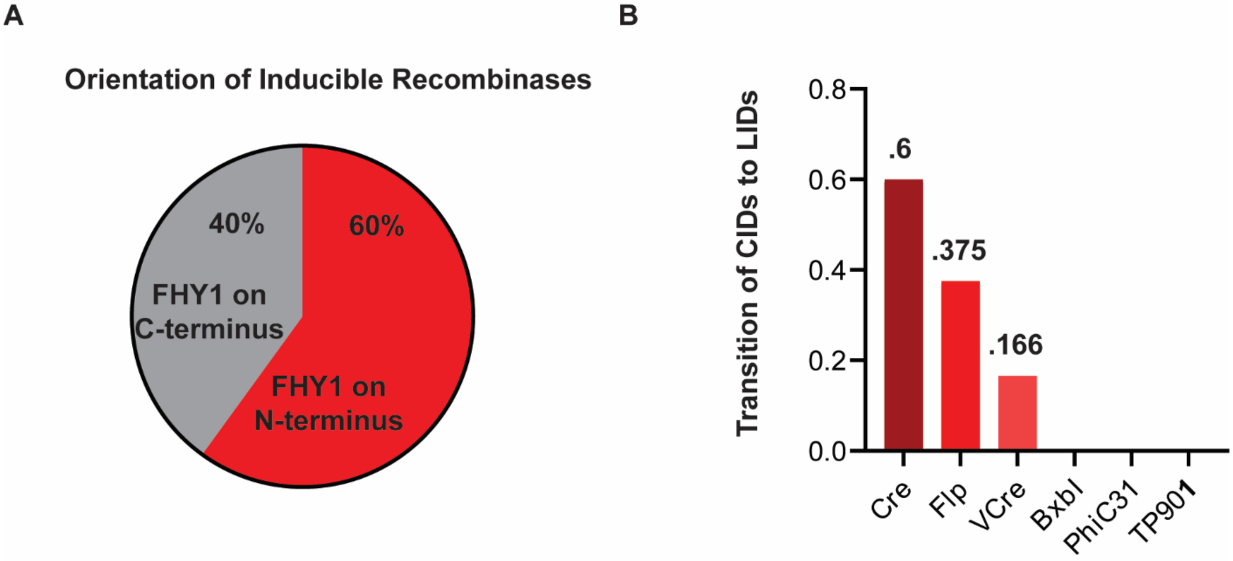


**Supplementary Figure 8. Trends from multidimensional screen**

(A) Percentage of inducible split sites with FHY1 on either the N-terminus or C-terminus of split recombinase. (B) Fraction of recombinases with fold change over 8, which we defined as the threshold for inducibility.

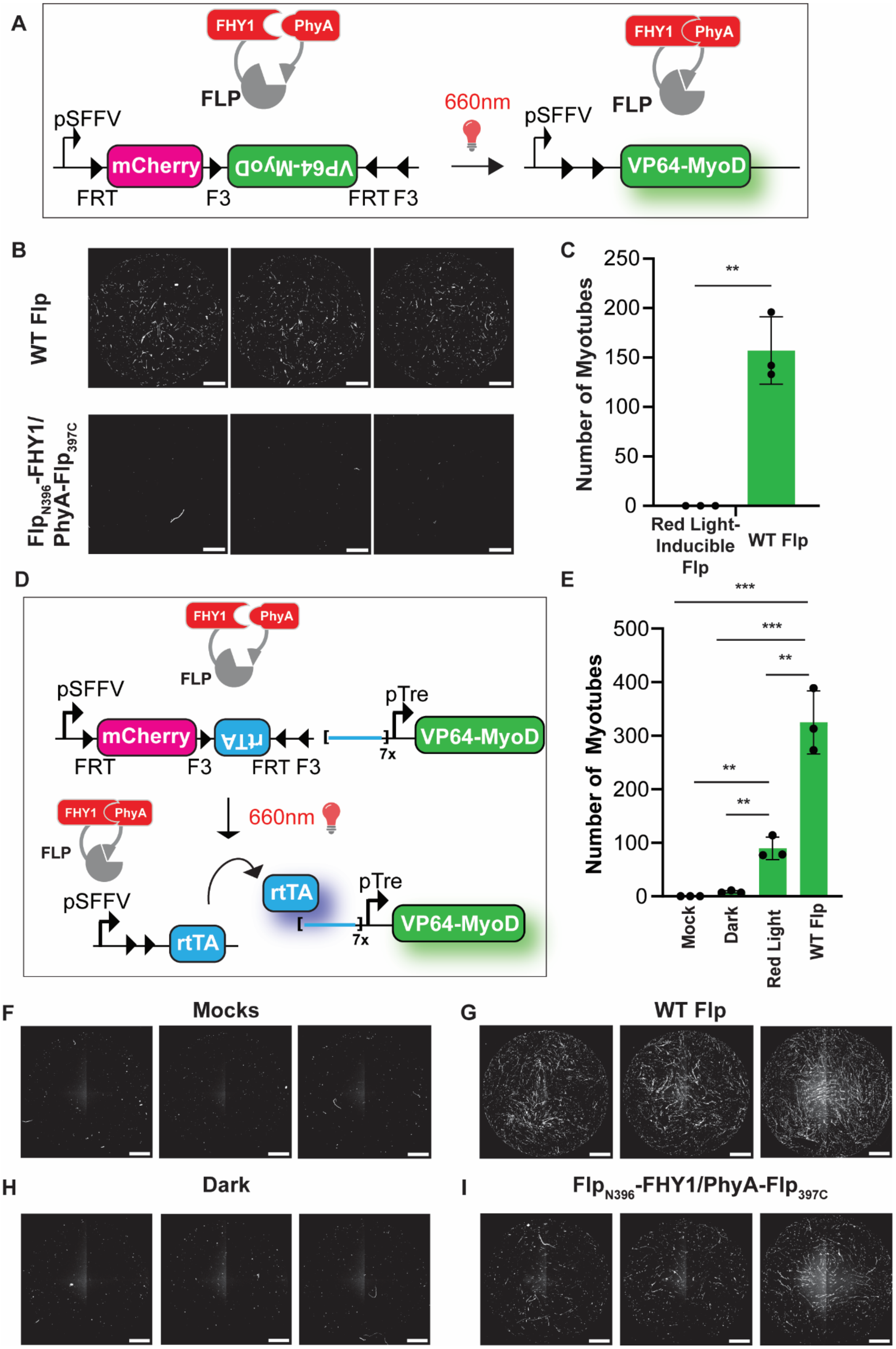
A
FHY1
PhyA
FLP
pSFFV
mCherry
VP64-MyoD
FRT
F3
FRT F3
660nm
B
WT Flp
FlpN396-FHY1/PhyA-Flp397C
C
Number of Myotubes
250
200
150
100
50
0
**
Red Light-Inducible Flp
WT Flp
D
rtTA
pTre
7x
E
500
400
300
200
100
***
**
Mock
Dark
Red Light
F
Mocks
G
H
Dark
I

**Supplementary Figure 9. Original MyoD circuit designs**

(A) Simplified non-amplifying circuit design was a FLEx switch where a Flp recombinase excises mCherry and flips on MyoD. (B) Images of anti-MHC immunostained $C3H_{myod}$ cells, with either transfected WT Flp (top panel) or $Flp_{N396}$-FHY1/PhyA-$Flp_{397C}$ (bottom panel). (C) Number of myotubes counted in each replicate. P-values were calculated by two-tailed unpaired t-test for n=3, p=0.0013. (D) Amplifying circuit design was a FLEx switch where a Flp recombinase excises mCherry and flips on an rtTA sequence. rtTA then binds upstream of VP64-MyoD to induce myogenesis. (E) Number of myotubes counted for $C3H_{rtTA}$ cells with either transfected mock control, WT Flp, or $Flp_{N396}$-FHY1/PhyA-$Flp_{397C}$ in either the dark or under red light (n=3). P-values were calculated as two-tailed unpaired t-test. For **p<0.01, for *** p<0.001 and for **** p<0.0001. (F) Images of anti-MHC immunostained $C3H_{rtTA}$ cells with transfected mock control (G) WT Flp, (H) $Flp_{N396}$-FHY1/PhyA-$Flp_{397C}$ in the dark (I) $Flp_{N396}$-FHY1/PhyA-$Flp_{397C}$ under red light. Scale bars are 1 mm.

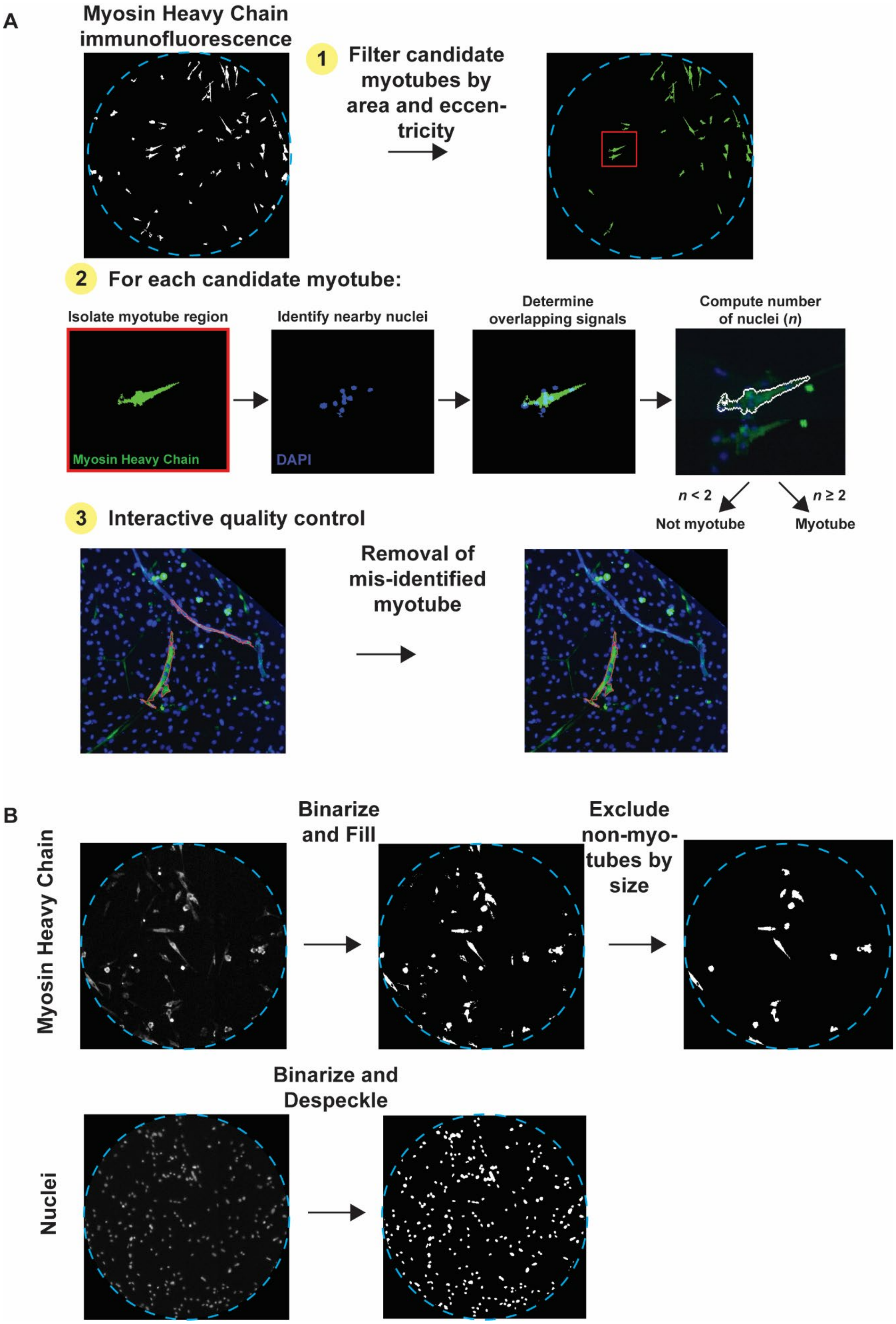

A
Myosin Heavy Chain immunofluorescence
1 Filter candidate myotubes by area and eccentricity
2 For each candidate myotube:
Isolate myotube region
Identify nearby nuclei
Determine overlapping signals
Compute number of nuclei (n)
Myosin Heavy Chain
DAPI
n < 2
n ≥ 2
Not myotube
Myotube
3 Interactive quality control
Removal of mis-identified myotube
B
Myosin Heavy Chain
Binarize and Fill
Exclude non-myotubes by size
Nuclei
Binarize and Despeckle

**Supplementary Figure 10. Classification of myotubes in myosin heavy chain (MHC) channel**

(A) For each structure in the MHC channel, our code identified nearby nuclei, overlayed them onto the MHC structure, counted the number of nuclei and annotated the image. This was repeated for each candidate. (B) Immunostained images are collected in 2 epifluorescence channels, 1 for MHC and 1 for nuclei. Images stained green for MHC are binarized and filled to boost signal and then processed to remove structures that are too small or circular. Images stained with DAPI for nuclei are binarized and despeckled to improve reads.

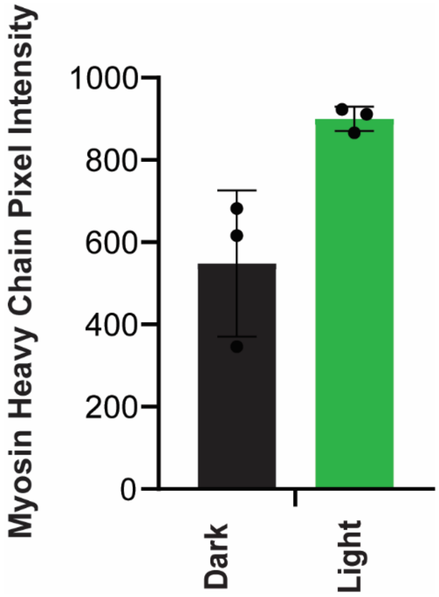


**Supplementary Figure 11. Myosin Heavy Chain (MHC) pixel intensity of illuminated vs. dark $C3H_{rtTA}$ cells**

Quantification of MHC chain pixel intensity in $C3H_{rtTA}$ cells transfected with $Flp_{N396}$-FHY1/PhyA-$Flp_{397C}$ and then either kept in the dark or red exposed to red light for 12 hours. Data represented as mean values ± SD (n=3).

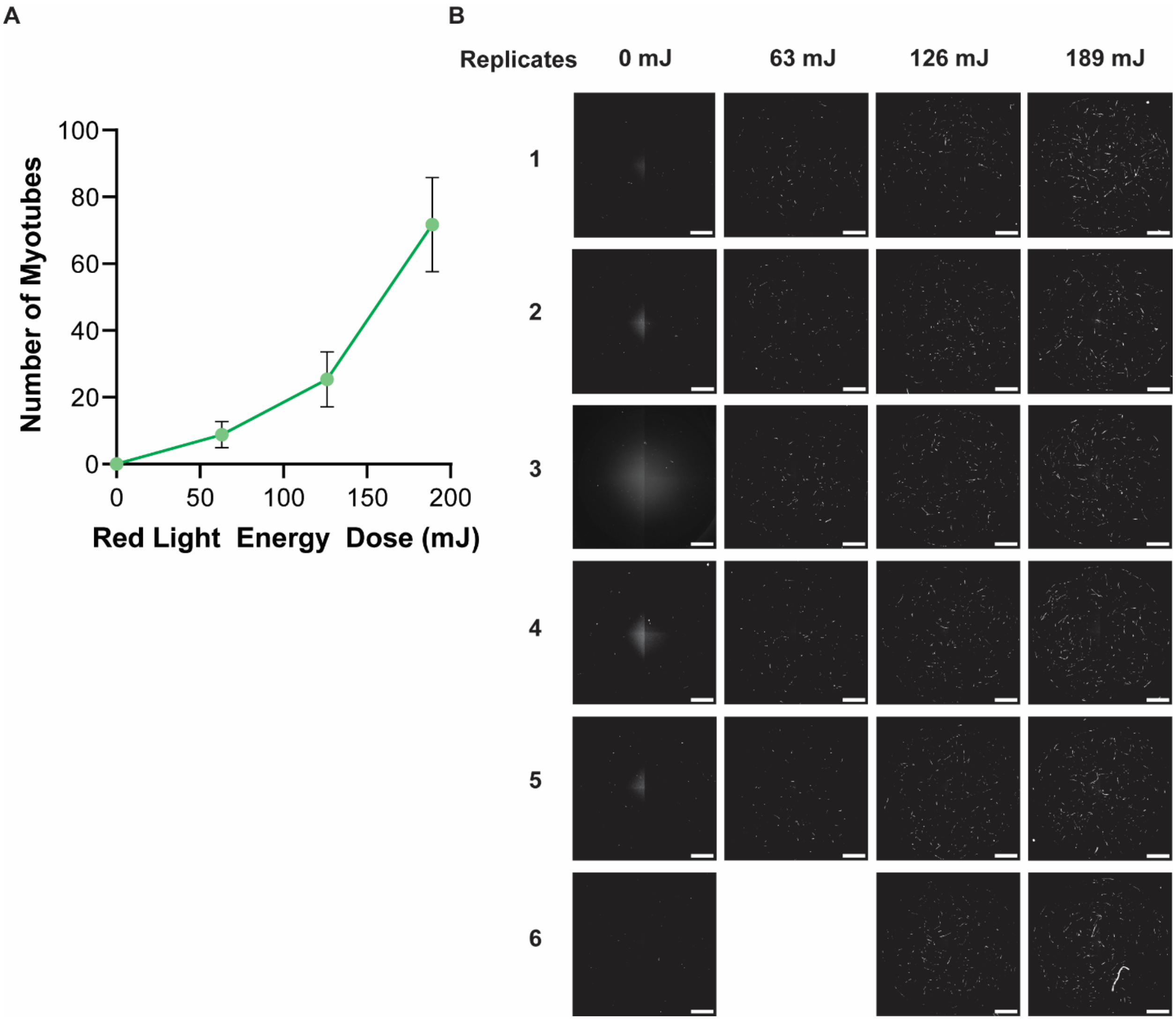


**Supplementary Figure 12. Titration of energy dose to induce myogenesis**

(A) Number of myotubes for $C3H_{tTA}$ cells (n=5 for 63 mJ condition and n=6 for others). (B) Epifluorescence images of MHC channel. All scale bars are 1 mm.

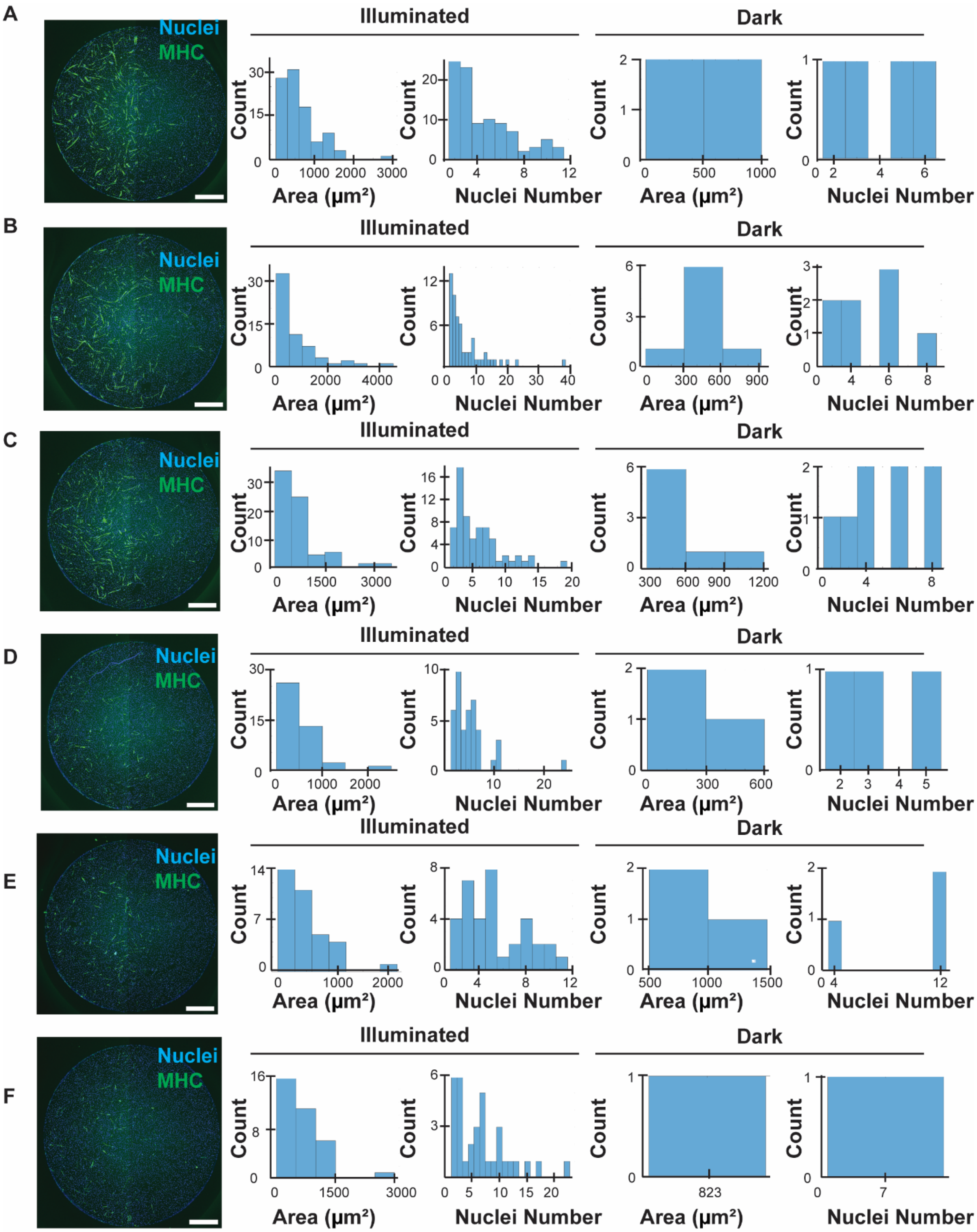


**Supplementary Figure 13. Histograms of area and nuclei number for red light patterned wells**

(A-F) Replicates of $C3H_{tTA}$ cells. From left to right: an epifluorescence image stained with DAPI (blue) and anti-MHC (green). Then, the respective area and nuclei number histograms are shown for the illuminated half of the well and the dark half. All scale bars are 1 mm.

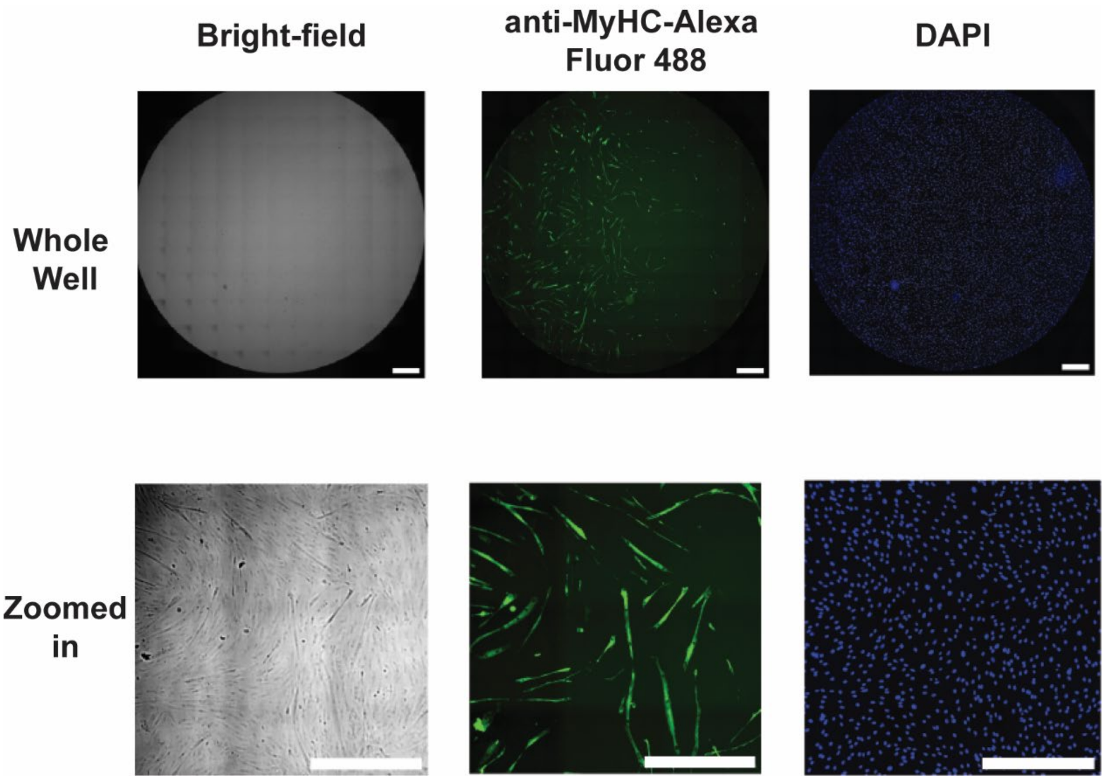


**Supplementary Figure 14. Brightfield and Epifluorescence images of red light-inducible myotubes**

Brightfield and epifluorescence images of $C3H_{tTA}$ cells. All scale bars are 0.5 mm.

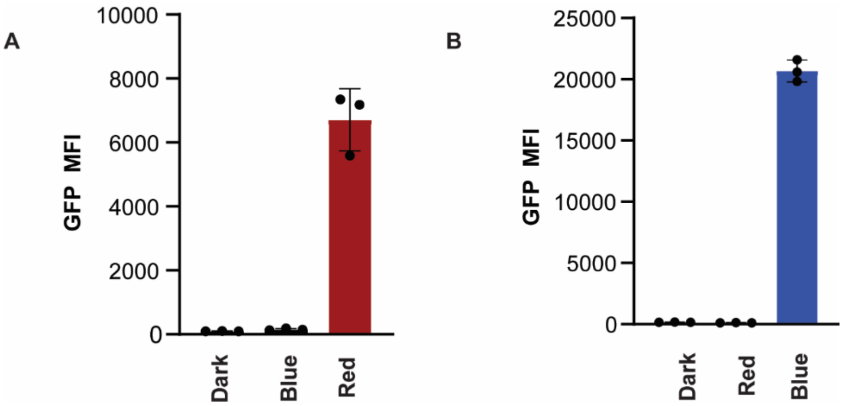


**Supplementary Figure 15. Orthogonality of red and blue light-inducible recombinases**

(A) $Flp_{N396}$-FHY1/PhyA-$Flp_{397C}$ transfected into HEK cells with GFP reporter and either red light, blue light, or no illumination. (B) $Cre_{N251}$-nMag/pMag-$Cre_{252C}$ transfected into HEK cells with a GFP reporter and either red light, blue light, or no illumination. Data represented as mean values ± SD (n=3).

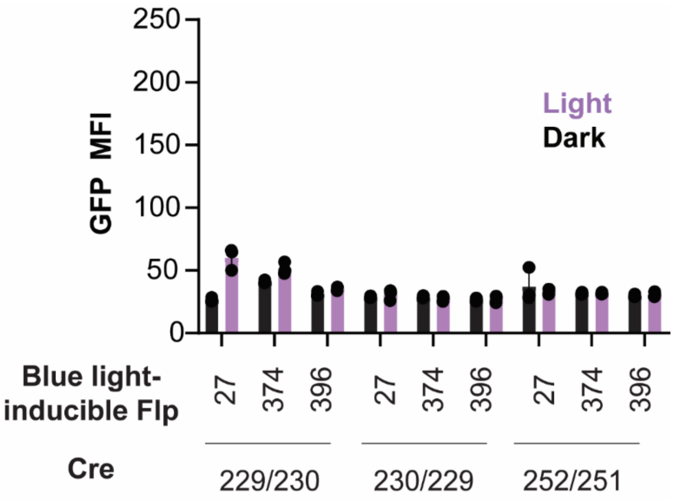


**Supplementary Figure 16. Characterization of multiplexed recombinases transfected with AND gate**

3 red light-inducible Cre recombinases and 3 blue light-inducible Flp recombinases were transfected into HEK cells with the AND gate circuit shown in Figure 3a. In the presence of red and blue light, these pairs did not increase GFP reporter expression. Data represented as mean values ± SD (n=3).

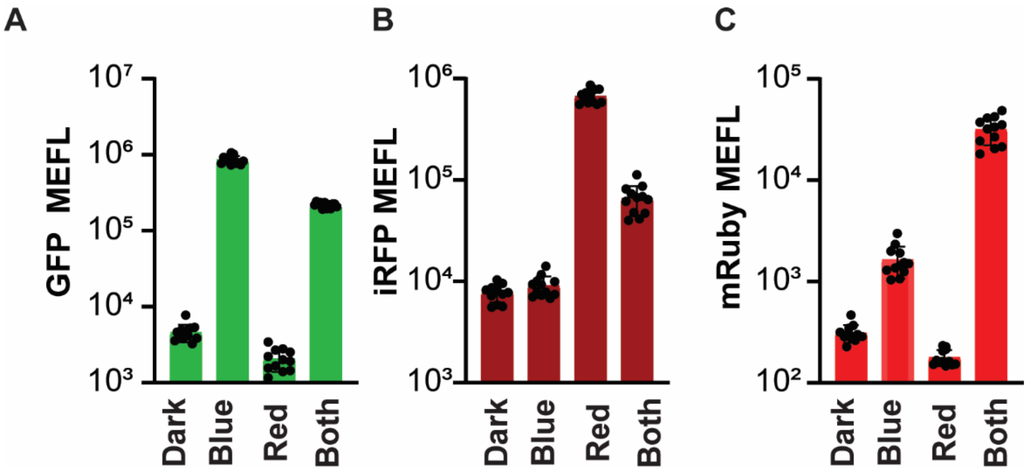


**Supplementary Figure 17. BLADE Circuit shown with a logscale**

$Flp_{N396}$-FHY1/PhyA-$Flp_{397C}$ and $Cre_{N251}$-nMag/pMag-$Cre_{252C}$ transfected into HEK cells with BLADE (same data as Fig. 3d shown on a logscale). MEFL shown for (A) GFP (BF) iRFP (C) mRuby in the dark, under blue illumination, red illumination, or both red and blue illumination. Data represented as mean equivalent fluorochrome values ± SD (n=12).

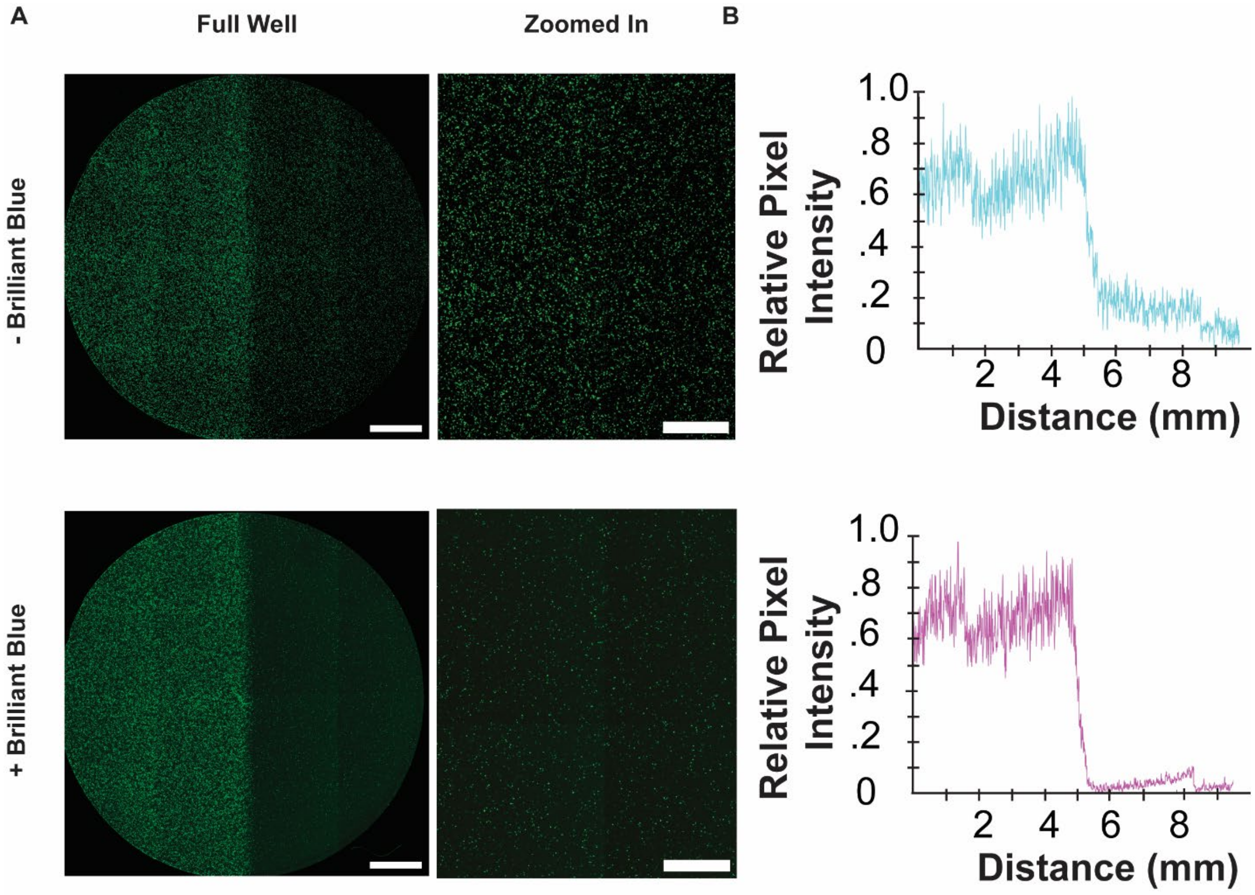


**Supplementary Figure 18. Effect of Brilliant Blue on cell patterning**

(A) $Flp_{N396}$-FHY1/PhyA-$Flp_{397C}$ and GFP reporter transfected into HEK cells. Epifluorescence image of GFP reporter expression in media with and without Brilliant Blue. Scale bars are 2 mm. (B) Corresponding intensity plot curve for image taken for each image.

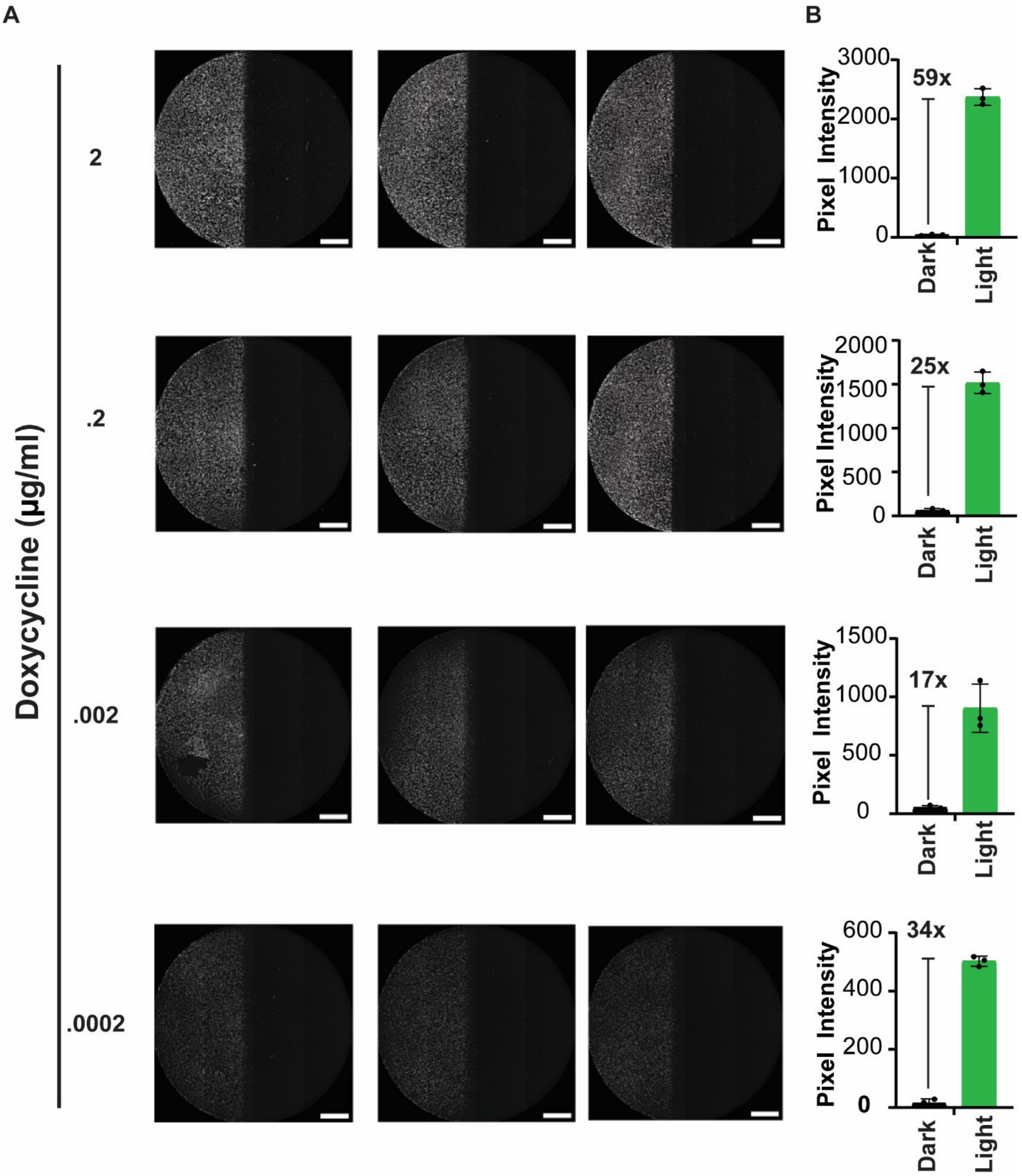


**Supplementary Figure 19. Titration of doxycycline with transfected red light-inducible Flp**

(A) Triplicate wells of HEK cells transfected with $Flp_{N396}$-FHY1/PhyA-$Flp_{397C}$ and illuminated with red light on the left half of the well. Scale bars are 2 mm. (B) Quantification of pixel intensity. Data represented as mean values ± SD (n=3).

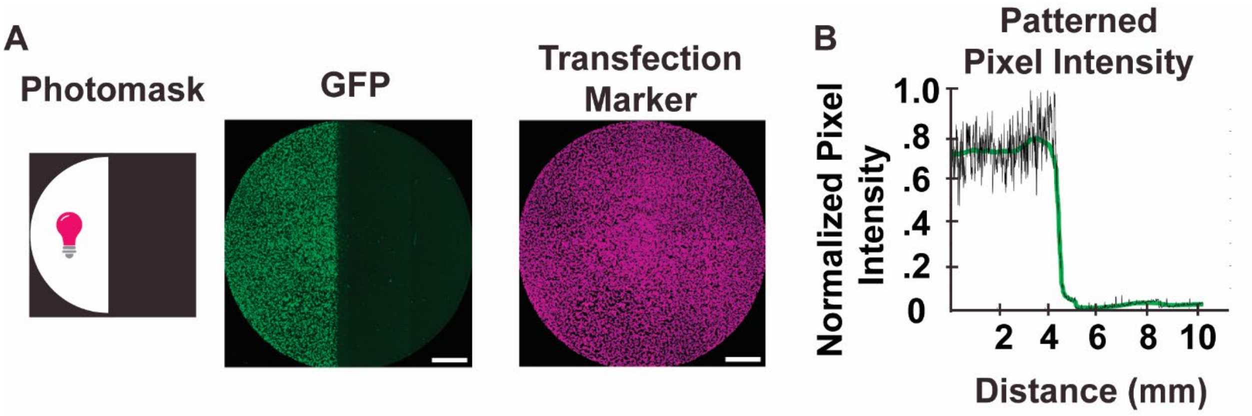


**Supplementary Figure 20. Red light-inducible Flp patterning**

(A) HEK cells transfected with $Flp_{N396}$-FHY1/PhyA-$Flp_{397C}$ and GFP reporter illuminated with red light on left half of the well. Epifluorescence channels are shown for GFP reporter and transfection marker. Scale bars are 2 mm. (B) Corresponding pixel intensity plot of GFP output.

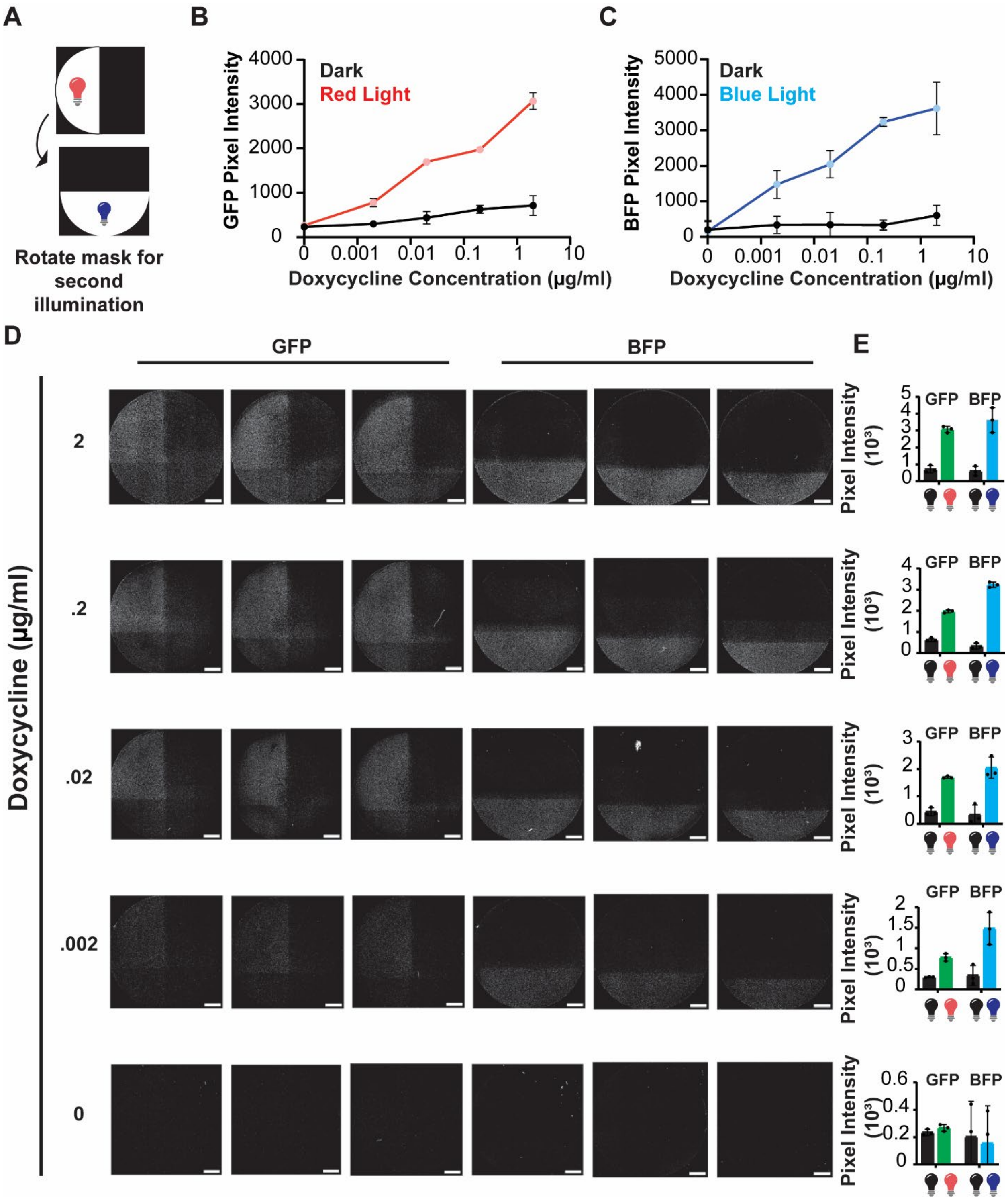


**Supplementary Figure 21. Doxycycline concentration of multiplexed recombinases**

(A) Schematic of photomask orientation for HEK cells transfected with $Flp_{N396}$-FHY1/PhyA-$Flp_{397C}$ and $Cre_{N251}$-nMag/pMag-$Cre_{252C}$. The left half of the well was illuminated with red light and the bottom half with blue light. (B) Graph of doxycycline titration and resulting GFP pixel intensity from red light-inducible Flp. (C) Graph of doxycycline titration and resulting BFP pixel intensity from blue light-inducible Cre. (D) Epifluorescence images are collected in either the GFP or BFP epifluorescence channel. (E) Bar chart of pixel intensity. Data represented as mean values ± SD (n=3). Scale bars are 2 mm.

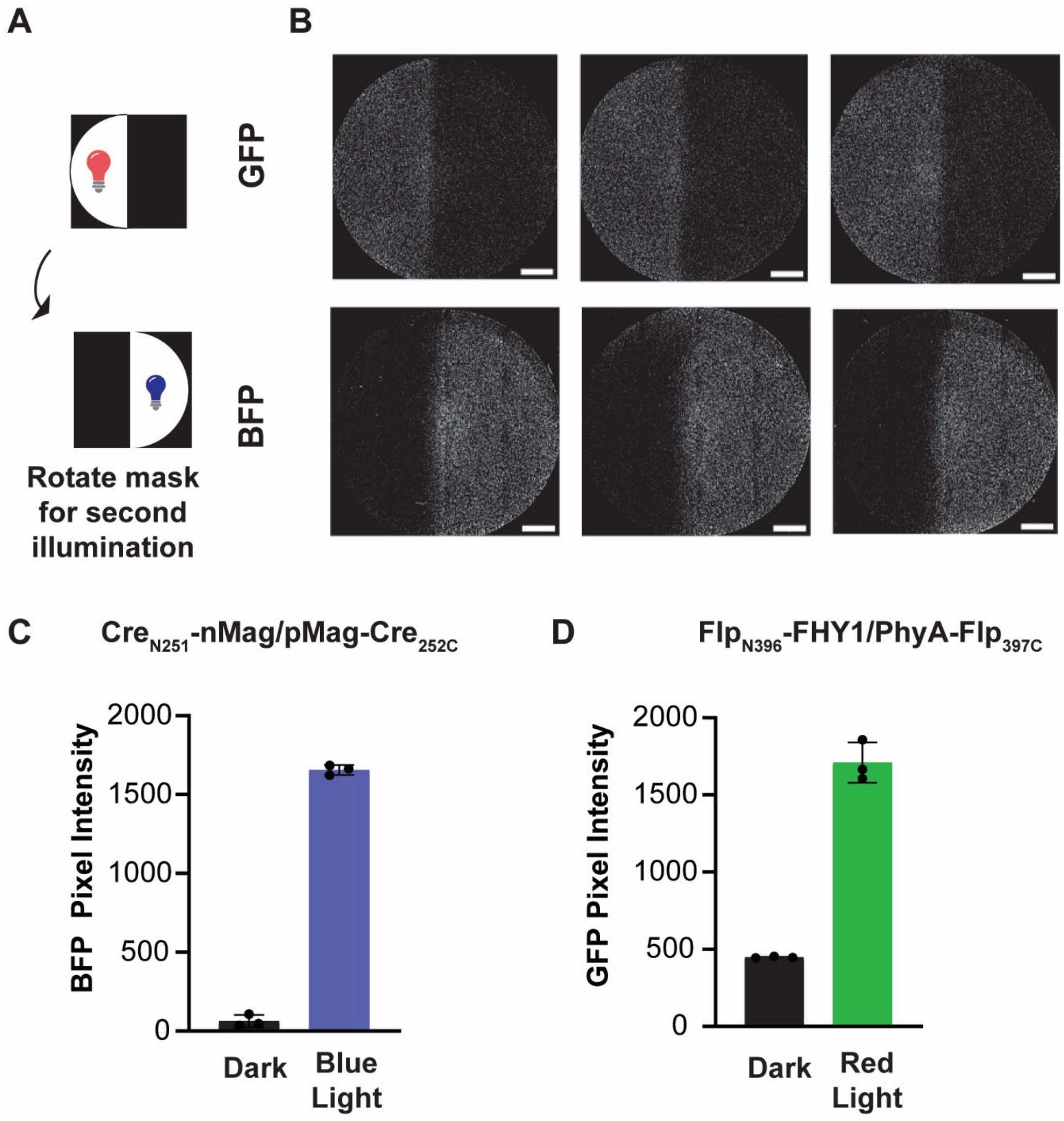


**Supplementary Figure 22. Red and blue light-inducible Flp and Cre patterned pixel intensity**

(A) Schematic of photomask orientation for HEK cells transfected with $Flp_{N396}$-FHY1/PhyA-$Flp_{397C}$ and $Cre_{N251}$-nMag/pMag-$Cre_{252C}$. The left half of the well was illuminated with red light and the right half of the well with blue light. (B) Epifluorescence images of either GFP or BFP. (C) Graph of BFP pixel intensity (n=3) (D) Graph of GFP pixel intensity. Data represented as mean values ± SD (n=3). Scale bars are 2 mm.

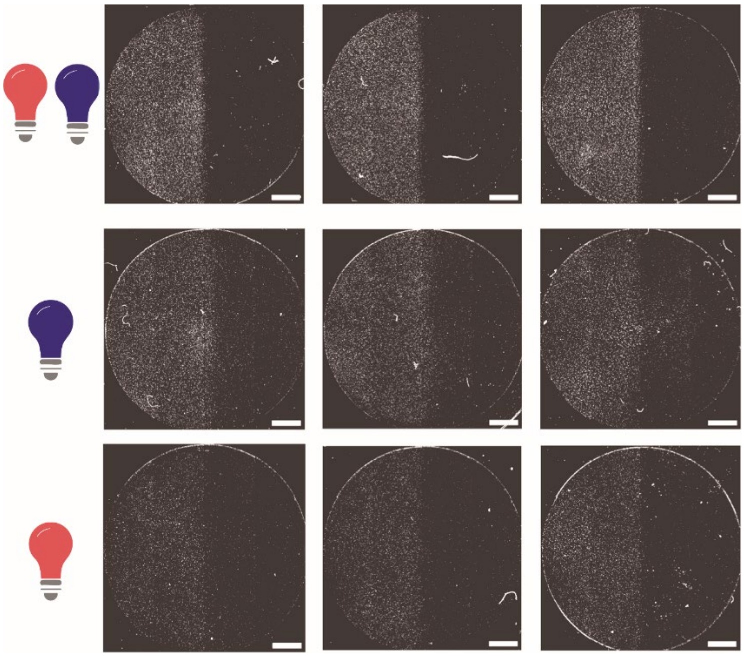

**Supplementary Figure 23. Triplicates for AND gate patterning**

Epifluorescence images of HEK cells transfected with $Flp_{N396}$-FHY1/PhyA-$Flp_{397C}$, $Cre_{N229}$-nMag/pMag-$Cre_{230C}$ with an AND gate reporter. Scale bars are 2 mm.

| Plasmid Name | Composition | Figure |
|---|---|---|
| CT361 | CT361_pCAG-PV1-DeltaPhyA-L1-iCre-106C-NLS-BGHpA | Figure 1 |
| CT362 | CT362_pCAG-PV1-DeltaPhyA-L1-iCre-230C-NLS-BGHpA | Figure 1 |
| CT363 | CT363_pCAG-PV1-DeltaPhyA-L1-iCre-252C-NLS-BGHpA | Figure 1 |
| CT364 | CT364_pCAG-PV1-DeltaPhyA-L1-iCre-257C-NLS-BGHpA | Figure 1 |
| CT365 | CT365_pCAG-PV1-DeltaPhyA-L1-iCre-271C-NLS-BGHpA | Figure 1 |
| CT366 | CT366_pCAG-PV1-iCre-N105-L1_FHY1-NLS-BGHpA | Figure 1 |
| CT367 | CT367_pCAG-PV1-iCre-N229-L1_FHY1-NLS-BGHpA | Figure 1 |
| CT368 | CT368_pCAG-PV1-iCre-N251-L1_FHY1-NLS-BGHpA | Figure 1 |
| CT369 | CT369_pCAG-PV1-iCre-N256-L1_FHY1-NLS-BGHpA | Figure 1 |
| CT370 | CT370_pCAG-PV1-iCre-N270-L1_FHY1-NLS-BGHpA | Figure 1 |
| CT351 | CT351_pCAG-PV1-iCre-N105-L1_Aff6-NLS-BGHpA | Figure 1 |
| CT352 | CT352_pCAG-PV1-iCre-N229-L1_Aff6-NLS-BGHpA | Figure 1 |
| CT353 | CT353_pCAG-PV1-iCre-N251-L1_Aff6-NLS-BGHpA | Figure 1 |
| CT354 | CT354_pCAG-PV1-iCre-N256-L1_Aff6-NLS-BGHpA | Figure 1 |
| CT355 | CT355_pCAG-PV1-iCreN270-L1-Aff6-NLS-BGHpA | Figure 1 |
| CT356 | CT356_pCAG-PV1-(FL)DrBphP-L1-iCre-106C-NLS-BGHpA | Figure 1 |
| CT357 | CT357_pCAG-PV1-(FL)DrBphP-L1-iCre-230C-NLS-BGHpA | Figure 1 |
| CT358 | CT358_pCAG-PV1-(FL)DrBphP-L1-iCre-252C-NLS-BGHpA | Figure 1 |
| CT359 | CT359_pCAG-PV1-(FL)DrBphP-L1-iCre-257C-NLS-BGHpA | Figure 1 |
| CT360 | CT360_pCAG-PV1-(FL)DrBphP-L1-iCre-271C-NLS-BGHpA | Figure 1 |
| CT438 | CT438-pCAG-PV1-FlpO-N27-L1-PhyA-NLS-BGHpA | Figure 1 |
| CT439 | CT439-pCAG-PV1-PhyA-L1-FlpO-28C-NLS-BGHpA | Figure 1 |

| | | |
|---|---|---|
| CT440 | CT440-pCAG-PV1-FlpO-N168-L1-PhyA-NLS-BGHpA | Figure 1 |
| CT441 | CT441-pCAG-PV1-PhyA-L1-FlpO-169C-NLS-BGHpA | Figure 1 |
| CT442 | CT442-pCAG-PV1-FlpO-N374-L1-PhyA-NLS-BGHpA | Figure 1 |
| CT443 | CT443-pCAG-PV1-PhyA-L1-FlpO-375C-NLS-BGHpA | Figure 1 |
| CT444 | CT444-pCAG-PV1-FlpO-N396-L1-PhyA-NLS-BGHpA | Figure 1 |
| CT445 | CT445-pCAG-PV1-PhyA-L1-FlpO-397C-NLS-BGHpA | Figure 1 |
| CT446 | CT446-pCAG-PV1-FlpO-N27-L1-FHY1-NLS-BGHpA | Figure 1 |
| CT447 | CT447-pCAG-PV1-FHY1-L1-FlpO-28C-NLS-BGHpA | Figure 1 |
| CT448 | CT448-pCAG-PV1-FlpO-N168-L1-FHY1-NLS-BGHpA | Figure 1 |
| CT449 | CT449-pCAG-PV1-FHY1-L1-FlpO-169C-NLS-BGHpA | Figure 1 |
| CT450 | CT450-pCAG-PV1-FlpO-N374-L1-FHY1-NLS-BGHpA | Figure 1 |
| CT451 | CT451-pCAG-PV1-FHY1-L1-FlpO-375C-NLS-BGHpA | Figure 1 |
| CT452 | CT452-pCAG-PV1-FlpO-N396-L1-FHY1-NLS-BGHpA | Figure 1 |
| CT453 | CT453-pCAG-PV1-FHY1-L1-FlpO-397C-NLS-BGHpA | Figure 1 |
| CT460 | CT460-pCAG-PV1-PhiC31-N233-L1-PhyA-NLS-BGHpA | Figure 1 |
| CT461 | CT461-pCAG-PV1-PhyA-L1-PhiC31-234C-NLS-BGHpA | Figure 1 |
| CT462 | CT462-pCAG-PV1-PhiC31-N396-L1-PhyA-NLS-BGHpA | Figure 1 |
| CT463 | CT463-pCAG-PV1-PhyA-L1-PhiC31-397C-NLS-BGHpA | Figure 1 |
| CT464 | CT464-pCAG-PV1-PhiC31-N428-L1-PhyA-NLS-BGHpA | Figure 1 |
| CT465 | CT465-pCAG-PV1-PhyA-L1-PhiC31-429C-NLS-BGHpA | Figure 1 |
| CT466 | CT466-pCAG-PV1-PhiC31-N571-L1-PhyA-NLS-BGHpA | Figure 1 |
| CT467 | CT467-pCAG-PV1-PhyA-L1-PhiC31-572C-NLS-BGHpA | Figure 1 |
| CT468 | CT468-pCAG-PV1-PhiC31-N233-L1-FHY1-NLS-BGHpA | Figure 1 |

| | | |
|---|---|---|
| CT469 | CT469-pCAG-PV1-FHY1-L1-PhiC31-234C-NLS-BGHpA | Figure 1 |
| CT470 | CT470-pCAG-PV1-PhiC31-N396-L1-FHY1-NLS-BGHpA | Figure 1 |
| CT471 | CT471-pCAG-PV1-FHY1-L1-PhiC31-397C-NLS-BGHpA | Figure 1 |
| CT472 | CT472-pCAG-PV1-PhiC31-N428-L1-FHY1-NLS-BGHpA | Figure 1 |
| CT473 | CT473-pCAG-PV1-FHY1-L1-PhiC31-429C-NLS-BGHpA | Figure 1 |
| CT474 | CT474-pCAG-PV1-PhiC31-N571-L1-FHY1-NLS-BGHpA | Figure 1 |
| CT475 | CT475-pCAG-PV1-FHY1-L1-PhiC31-572C-NLS-BGHpA | Figure 1 |
| CT508 | CT508_pCAG-PV1-BX-N197-L1-PhyA-NLS-BGHpA | Figure 1 |
| CT509 | CT509_pCAG-PV1-BX-N197-L1-FHY1-NLS-BGHpA | Figure 1 |
| CT510 | CT510_pCAG-PV1-PhyA-L1-BX-198C-NLS-BGHpA | Figure 1 |
| CT511 | CT511_pCAG-PV1-FHY1-L1-BX-198C-NLS-BGHpA | Figure 1 |
| CT512 | CT512_pCAG-PV1-BX-N468-L1-PhyA-NLS-BGHpA | Figure 1 |
| CT513 | CT513_pCAG-PV1-BX-N468-L1-FHY1-NLS-BGHpA | Figure 1 |
| CT514 | CT514_pCAG-PV1-PhyA-L1-BX-469C-NLS-BGHpA | Figure 1 |
| CT515 | CT515_pCAG-PV1-FHY1-L1-BX-469C-NLS-BGHpA | Figure 1 |
| CT516 | CT516_pCAG-PV1-TP901-N323-L1-PhyA-NLS-BGHpA | Figure 1 |
| CT517 | CT517_pCAG-PV1-TP901-N323-L1-FHY1-NLS-BGHpA | Figure 1 |
| CT518 | CT518_pCAG-PV1-PhyA-L1-TP901-324C-NLS-BGHpA | Figure 1 |
| CT519 | CT519_pCAG-PV1-FHY1-L1-TP901-324C-NLS-BGHpA | Figure 1 |
| CT520 | CT520_pCAG-PV1-TP901-N326-L1-PhyA-NLS-BGHpA | Figure 1 |
| CT521 | CT521_pCAG-PV1-TP901-N326-L1-FHY1-NLS-BGHpA | Figure 1 |
| CT522 | CT522_pCAG-PV1-PhyA-L1-TP901-327C-NLS-BGHpA | Figure 1 |
| CT523 | CT523_pCAG-PV1-FHY1-L1-TP901-327C-NLS-BGHpA | Figure 1 |

| | | |
|---|---|---|
| CT524 | CT524_pCAG-PV1-VCre-N172-L1-PhyA-NLS-BGHpA | Figure 1 |
| CT525 | CT525_pCAG-PV1-VCre-N172-L1-FHY1-NLS-BGHpA | Figure 1 |
| CT526 | CT526_pCAG-PV1-PhyA-L1-VCre-173C-NLS-BGHpA | Figure 1 |
| CT527 | CT527_pCAG-PV1-FHY1-L1-VCre-173C-NLS-BGHpA | Figure 1 |
| CT528 | CT528_pCAG-PV1-VCre-N269-L1-PhyA-NLS-BGHpA | Figure 1 |
| CT529 | CT529_pCAG-PV1-VCre-N269-L1-FHY1-NLS-BGHpA | Figure 1 |
| CT530 | CT530_pCAG-PV1-PhyA-L1-VCre-270C-NLS-BGHpA | Figure 1 |
| CT531 | CT531_pCAG-PV1-FHY1-L1-VCre-270C-NLS-BGHpA | Figure 1 |
| CT532 | CT532_pCAG-PV1-VCre-N277-L1-PhyA-NLS-BGHpA | Figure 1 |
| CT533 | CT533_pCAG-PV1-VCre-N277-L1-FHY1-NLS-BGHpA | Figure 1 |
| CT534 | CT534_pCAG-PV1-PhyA-L1-VCre-278C-NLS-BGHpA | Figure 1 |
| CT535 | CT535-pCAG-PV1-FHY1-L1-VCre-278C-NLS-BGHpA | Figure 1 |
| CT620 | CT620_pCAG-PV1-DrBphP-L1-iCre-106C-NLS-BGHpA | Figure 1 |
| CT621 | CT621_pCAG-PV1-DrBphP-L1-iCre-230C-NLS-BGHpA | Figure 1 |
| CT622 | CT622_pCAG-PV1-DrBphP-L1-iCre-252C-NLS-BGHpA | Figure 1 |
| CT623 | CT623_pCAG-PV1-DrBphP-L1-iCre-257C-NLS-BGHpA | Figure 1 |
| CT624 | CT624_pCAG-PV1-DrBphP-L1-iCre-271C-NLS-BGHpA | Figure 1 |
| CT625 | CT625_pCAG-PV1-iCre-N105-L1_LDB3-NLS-BGHpA | Figure 1 |
| CT626 | CT626_pCAG-PV1-iCre-N229-L1_LDB3-NLS-BGHpA | Figure 1 |
| CT627 | CT627_pCAG-PV1-iCre-N251-L1_LDB3-NLS-BGHpA | Figure 1 |
| CT628 | CT628_pCAG-PV1-iCre-N256-L1_LDB3-NLS-BGHpA | Figure 1 |
| CT629 | CT629_pCAG-PV1-iCre-N270-L1_LDB3-NLS-BGHpA | Figure 1 |
| CT610 | CT610_pCAG-PV1-FHY1-L1-iCre-106C-NLS-BGHpA | Figure 1 |

| | | |
|---|---|---|
| CT611 | CT611_pCAG-PV1-FHY1-L1-iCre-230C-NLS-BGHpA | Figure 1 |
| CT612 | CT612_pCAG-PV1-FHY1-L1-iCre-252C-NLS-BGHpA | Figure 1 |
| CT613 | CT613_pCAG-PV1-FHY1-L1-iCre-257C-NLS-BGHpA | Figure 1 |
| CT614 | CT614_pCAG-PV1-FHY1-L1-iCre-271C-NLS-BGHpA | Figure 1 |
| CT615 | CT615_pCAG-PV1-iCre-N105-L1_PhyA-NLS-BGHpA | Figure 1 |
| CT616 | CT616_pCAG-PV1-iCre-N229-L1_PhyA-NLS-BGHpA | Figure 1 |
| CT617 | CT617_pCAG-PV1-iCre-N251-L1_PhyA-NLS-BGHpA | Figure 1 |
| CT618 | CT618_pCAG-PV1-iCre-N256-L1_PhyA-NLS-BGHpA | Figure 1 |
| CT619 | CT619_pCAG-PV1-iCre-N270-L1_PhyA-NLS-BGHpA | Figure 1 |
| CT750 | CT750_pSFFV-frt-mCh-f3-rtTa-frt-f3 | Figure 2 |
| CT751 | CT751_pTRE3G-VP64-MyoD-T2A-iRFP-frt-f3 | Figure 2 |
| CT754 | CT754_pHR_pSFFV-frt-mCh-f3-tTa-frt-f3 | Figure 2 |
| CT769 | CT769_pHR-EF1a-PhyA-Flp397C-P2A-FlpN396-FHY1 | Figure 2 |
| BW336 | BW336_pCAG-frt-frt-loxp-loxp-GFP | Figure 3 |
| CT691 | CT691-BLADE DECODER | Figure 3 |
| CT752 | CT752_pCAG-FRT-3xpolyA-FRT-GFP | Figure 4 |
| CT641 | CT641_pCAG-FRT-Lox2272-polyA-Lox2272-GFP-FRT-loxP-LssmOrange-loxP-mRuby2 | Figure 4 |
| IK216 | IK216_pTre-iCre-N229-L1-pMag-NLS | Figure 4 |
| IK331 | IK331_pTre-iCre-N229-L1-pMag-NLS | Figure 4 |
| IK332 | IK332_pTre-iCre-nMag-L1-230C-NLS | Figure 4 |
| IK349 | IK349_pTre3G-PhyA-L1-Flp0_397C-NLS | Figure 4 |
| IK350 | IK350_pTre3G-FRT-FlpO-N396-L1-FHY1-NLS | Figure 4 |
| BW2144 | BW2144_PBP-pEF1a-rtTa | Figure 4 |

**Supplementary Table 1.** List of plasmids used in manuscript.

| Red Light Flp Split 1 | Red Light Flp Split 2 | Transfection Marker | Reporter |
|---|---|---|---|
| 31.6 ng | 31.6 ng | 7 ng | 31.6 ng |

**Supplementary Table 2.** Red light-inducible recombinase transfection ratios for flow cytometry experiments in 96 well plate.

| Red Light Flp Split 1 | Red Light Flp Split 2 | Blue Light Cre Split 1 | Blue Light Cre Split 2 | Transfection Marker | Reporter |
|---|---|---|---|---|---|
| 16.25 ng | 16.25 ng | 16.25 ng | 16.25 ng | 4 ng | 32.5 ng |

**Supplementary Table 3.** AND Gate transfection ratios for flow cytometry experiments in 96 well plate.

| Red Light Flp Split 1 | Red Light Flp Split 2 | Blue Light Cre Split 1 | Blue Light Cre Split 2 | Transfection Marker | Blank | Reporter |
|---|---|---|---|---|---|---|
| 14.28 ng | 14.28 ng | 14.28 ng | 14.28 ng | 14.28 ng | 14.28 ng | 14.28 ng |

**Supplementary Table 4.** BLADE transfection ratios for flow cytometry experiments in 96 well plate.

| Red Light Flp Split 1 (IK349) | Red Light Flp Split 2 (IK350) | Transfection Marker (BW463) | Reporter (CT752) | rtTA |
|---|---|---|---|---|
| 1216.02 ng | 1216.02 ng | 1365 ng | 14486.94 ng | 1216.02 ng |

**Supplementary Table 5.** Red light-inducible recombinase transfection ratios for patterning experiments in T-75.

| Red Light Flp Split 1 (IK349) | Red Light Flp Split 2 (IK350) | Blue Light Cre Split 1 (IK331) | Blue Light Cre Split 2 (IK332) | Transfection Marker | Flp Reporter (CT752) | Cre Reporter (IK216) | rtTA |
|---|---|---|---|---|---|---|---|
| 1218.75 ng | 1218.75 ng | 1218.75 ng | 1218.75 ng | 1218.75 ng | 6093.75 ng | 6093.7 5ng | 1218.75 ng |

**Supplementary Table 6.** Red and blue light-inducible recombinase transfection ratios for patterning experiments with individual recombinase reporters in T-75.

| Red Light Flp Split 1 (IK349) | Red Light Flp Split 2 (IK350) | Blue Light Cre Split 1 (IK331) | Blue Light Cre Split 2 (IK332) | Transfection Marker | AND Gate Reporter (BW336) | rtTA |
|---|---|---|---|---|---|---|
| 1950 ng | 1950 ng | 1950 ng | 1950 ng | 1950 ng | 7800 ng | 1950 ng |

**Supplementary Table 7.** Red and blue light-inducible recombinase transfection ratios for patterning experiments with AND gate in T-75.
